# Epigenome and early selection determine the tumour-immune evolutionary trajectory of colorectal cancer

**DOI:** 10.1101/2024.02.12.579956

**Authors:** Eszter Lakatos, Vinaya Gunasri, Luis Zapata, Jacob Househam, Timon Heide, Nicholas Trahearn, Ottilie Swinyard, Luis Cisneros, Claire Lynn, Maximilian Mossner, Chris Kimberley, Inmaculada Spiteri, George D. Cresswell, Gerard Llibre-Palomar, Miriam Mitchison, Carlo C. Maley, Marnix Jansen, Manuel Rodriguez-Justo, John Bridgewater, Ann-Marie Baker, Andrea Sottoriva, Trevor A. Graham

## Abstract

Immune system control is a major hurdle that cancer evolution must circumvent. The relative timing and evolutionary dynamics of subclones that have escaped immune control remain incompletely characterized, and how immune-mediated selection shapes the epigenome has received little attention. Here, we infer the genome- and epigenome-driven evolutionary dynamics of tumour-immune coevolution within primary colorectal cancers (CRCs). We utilise our existing CRC multi-region multi-omic dataset that we supplement with high-resolution spatially-resolved neoantigen sequencing data and highly multiplexed imaging of the tumour microenvironment (TME). Analysis of somatic chromatin accessibility alterations (SCAAs) reveals frequent somatic loss of accessibility at antigen presenting genes, and that SCAAs contribute to silencing of neoantigens. We observe that strong immune escape and exclusion occur at the outset of CRC formation, and that within tumours, including at the microscopic level of individual tumour glands, additional immune escape alterations have negligible consequences for the immunophenotype of cancer cells. Further minor immuno-editing occurs during local invasion and is associated with TME reorganisation, but that evolutionary bottleneck is relatively weak. Collectively, we show that immune evasion in CRC follows a “Big Bang” evolutionary pattern, whereby genetic, epigenetic and TME-driven immune evasion acquired by the time of transformation defines subsequent cancer-immune evolution.

## INTRODUCTION

During cancer evolution, tumours are continuously shaped by interactions with their environment, especially by the ongoing “war” with the immune system. Neoantigens – novel peptides originating from cancer-specific mutations – elicit immune recognition. Cancers may carry tens to hundreds of neoantigen-candidate mutations, but tumours ultimately evade immune elimination via immune-editing (losing antigenic mutations), immune-escape (e.g. hampering antigen presentation), immune exclusion (manipulating the tumour microenvironment to limit immune presence) or a combination of these mechanisms^1^. Immune evasion is unlikely to be a binary on/off trait, but a continuous phenotype modulated by the strength of the contribution of each of the three factors. Immunotherapies aim at re-invigorating and re-engaging the immune system in its war against cancer, and therefore understanding the mechanisms underlying immune evasion is paramount for treatment success.

Colorectal cancers (CRCs) generally display a relatively active immune microenvironment with substantial tumour-infiltrating lymphocytes^2^. About 15% of CRCs are mismatch repair deficient (MMRd), which is associated with a higher neoantigen burden, enhanced immune presence^3,4^ and good response to immune checkpoint blockade therapies (ICBs), however, still up to 30% of MMRd CRCs do not respond^5^. Mismatch repair proficient (MMRp) CRCs harbour a lower mutation burden than MMRd CRCs and ICB treatments are ineffective^6^, indicating that immune evasion in MMRp tumours likely occurs through mechanisms alternative to those targeted by current ICB drugs. Nonetheless, immune infiltrate levels are known to be prognostic for CRC^7^ indicating a central role for the immune system in cancer evolution. Indeed, most MMRp cancers carry (multiple) putative clonal neoantigen mutations^8^, and while only a subset of these might be sufficiently presented^9^, they may still be capable of T-cell activation^10–12^.

Immune evasion through genetic means (e.g. mutations that reduce the immune system’s ability to engage with neoantigens) has been thoroughly investigated and found to be common in CRCs^4,13^. Our previous mathematical modelling found that escape is essential for MMRd CRC development, and is a crucial step in MMRp CRC formation when immune surveillance is stringent^4,8,14^. The role of the epigenome in modulating tumour antigenicity, however, has received less attention: somatic changes in chromatin organisation are known to contribute to the cancer immunophenotype, but their impact on immune escape and editing have not been completely assessed^15–17^. Promoter hypermethylation has been identified as mechanism of neoantigen silencing in lung cancer^18^. Analogously, closing chromatin may lead to repression of antigen expression, meaning that immune selection could strongly shape a cancer’s epigenome architecture.

Previous analyses relied on bulk genetic sequencing data from superficial tumour, and so were likely unable to detect subclonal and heterogeneous immune evolutionary dynamics that could be associated with disease spread. Further, the invasive margin – the cancer region in direct contact to host tissue – is likely a major determinant of the overall immune evasion/elimination of the cancer, but has rarely been studied directly because it is typically fixed for diagnostic purposes and unavailable for research^19,20^. In summary, spatial heterogeneity in immunoediting and escape within CRCs remains incompletely characterised.

Here, we firstly explore the interplay between immune evasion and the epigenome, highlighting a role for chromatin architecture in suppressing expression of neoantigens and antigen presenting machinery in CRC. We explore individual tumour gland-level characterisation of immune evasion (microenvironment restructuring and genetic immune escape), exploring the intra-tumour heterogeneity within and across different morphological contexts. We leverage our existing multi-region multi-omic sequencing (matched genome, transcriptome and chromatin accessibility profiling) of 495 single glands (representing 29 CRCs) from our previously published Evolutionary Predictions in Colorectal Cancer (EPICC) study^15,21^. These data are supplemented with newly generated data that combines targeted neoantigen sequencing and highly multiplexed tumour microenvironment profiling of 82 micro-biopsies representing distinct tumour-associated regions from a subset of 11 EPICC cases, including lymph node metastases and distant normal mucosa.

## RESULTS

### Multi-modal immune analysis of CRC micro-biopsies

We analysed the immune landscape of 29 colorectal adenocarcinomas at single tumour gland resolution, using multi-omic (whole genome-sequencing (WGS), RNAseq and ATACseq) analysis of fresh frozen (FF) single-glands, coupled with high-depth panel sequencing (PS) and cyclic immunofluorescence (CyCIF^22^) imaging of matched formalin-fixed paraffin-embedded (FFPE) biopsies. The collection and processing of FF samples, part of our EPICC study, have been previously described in refs^15,21^ (Fig 1a). We also obtained diagnostic FFPE blocks from 11 patients who had MMRp stage III cancers with lymph node metastasis. We micro-dissected small clusters of glands representing the superficial tumour, invasive margin, and lymph node deposit (Fig 1a) and sequenced genomic regions associated with the immuno-peptidome^23^ (Methods). In a subset of these regions, we concurrently visualised 22 proteins that identified immune cell types or regulatory receptor expression (see Table S1) using CyCIF (Methods). In total, we assembled new and existing sequencing data from a total of 495 FF and 82 FFPE biopsies from 29 patients (median of 15 FF and 8 FFPE biopsies per patient) with concurrent CyCIF analysis of 8 patients.

**Figure 1.**
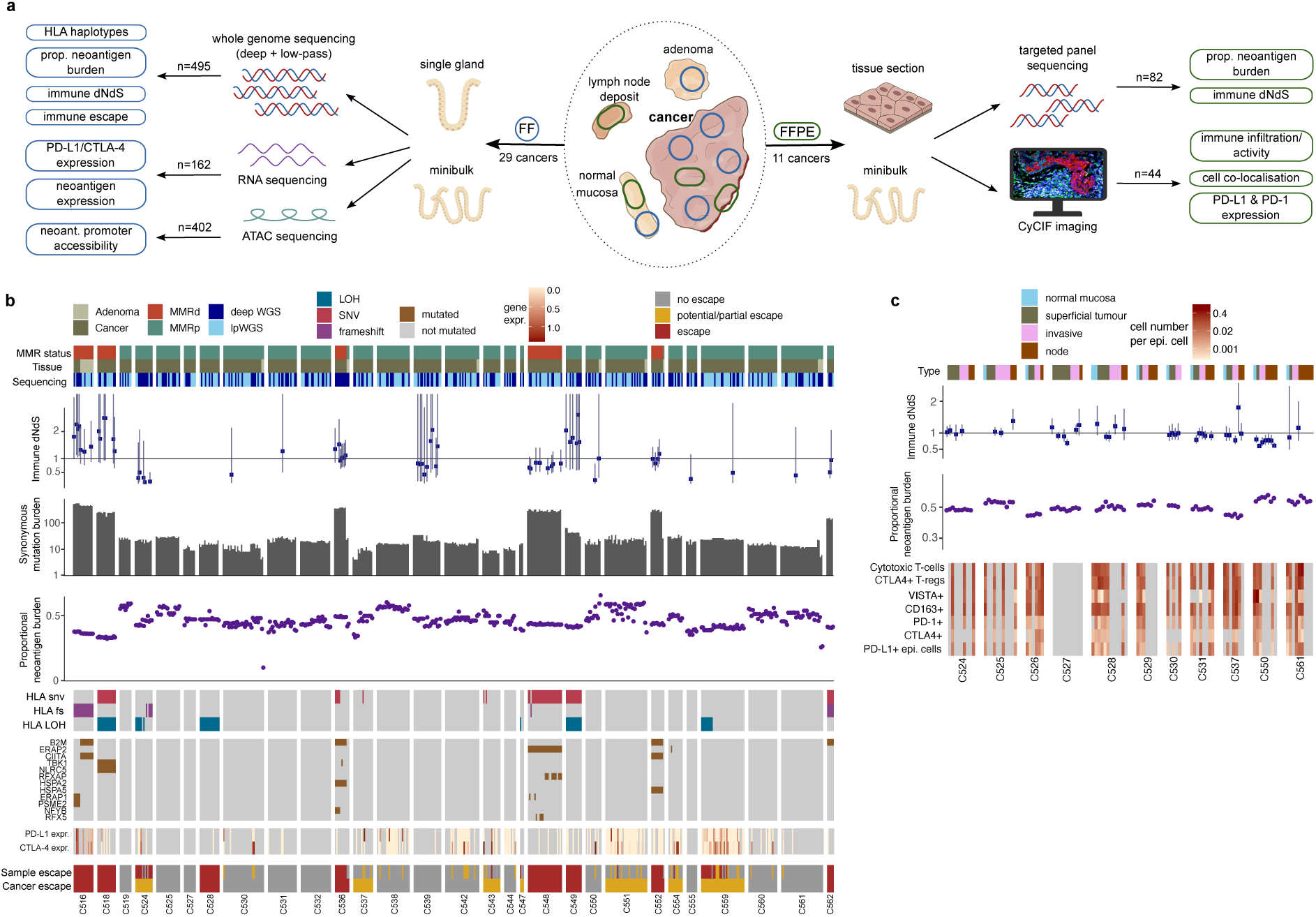
Multi-omic sequencing and multiplex imaging of the cancer-immune landscape of CRCs. (a) Overview of sample collection (middle), processing and analysis of FF (left) and FFPE (right) biopsies. (b) Neoantigen burden, synonymous mutation burden, immune dNdS and immune escape mutation overview of all FF-WGS samples. Error bars depict confidence intervals for immune dNdS estimates, truncated at 3.5. (c) Neoantigen burden, immune dNdS and immune infiltrate overview of all FFPE-PS samples.

In both whole genome- and panel-sequenced biopsies, we predicted the antigenicity of each somatic mutation using the NeoPredPipe pipeline^24^. We defined the (proportional) neoantigen burden of each biopsy as the number of mutations present that gave rise to at least one strong-binding (rank score <0.5), cancer-specific mutated peptide, normalised to the total burden of protein-changing mutations. For samples with sufficient mutations, we also estimated immune selection using the ratio of nonsynonymous to synonymous mutations in the immunopeptidome (immune dN/dS) using SOPRANO^23^ (Fig 1a), normalised to the expected ratio given the mutational spectra of all mutations.

In FF-WGS samples, we characterised candidate immune escape alterations: single nucleotide variants (SNVs), frameshifts and loss-of-heterozygosity (LOH) in HLA genes, high-impact alterations in other antigen presenting genes (APGs, see Methods for list) and high expression of PD-L1 or CTLA-4 (as measured by RNAseq). Samples with a LOH, frameshift, stop-gain alteration, or multiple SNVs in putative escape genes, or with over-expression of PD-L1 and CTLA-4 were labelled “escaped”, while samples with a single SNV or moderate PD-L1/CTLA-4 expression were labelled “potential escape”.

Neoantigen burden, immune dN/dS, genetic immune escape alterations and immune markers each showed high inter-patient but moderate intra-patient variability (Fig 1b,c). MMRd cancers (n=5) had a significantly higher overall mutation load than MMRp cancers (median synonymous burden 307 vs 18; median total neoantigen burden 292 vs 23, p<10^−16^) and all carried clonal high-impact immune escape alterations, associated with significant loss-of-function in antigen presentation through e.g. truncating frameshift mutations (Fig. 1b, Fig. S1). Immune escape alterations were detected in 14 of 29 cancers. They showed remarkable homogeneity (across multiple tumour regions) in 8/14 cancers and were subclonal in 6/14 cases (all MMRp, Fig. S1).

### Epigenetic and transcriptomic regulation of antigen presenting genes

We examined the role of the epigenome in enabling immune escape. We looked for somatic chromatin accessibility alterations (SCAAs) associated with immune escape at a list of predefined antigen presenting genes (APGs, see Methods) by examining the ATACseq data available from 25 CRCs. We detected a total of 45 SCAAs, all in promoter regions upstream of these APGs. 21/34 (62%) APGs had at least one SCAA and 9/25 (36%) of patients had at least one immune escape gene affected by a SCAA. Notably, 42 (93%) of the SCAAs were losses of accessibility, significantly different than expected based on the genome-wide distribution of SCAAs (p=0.025; see Methods). None of the SCAAs co-occurred with somatic mutations in those genes (Fig. 2a). This exclusivity could indicate that mutations are more likely to be found in expressed genes or suggest SCAAs as an alternative route to antigen presentation disruption. ERAP2 (SCAAloss in three cancers and three adenomas), for example, had considerably lower RNA expression in the affected cancers than in normal tissue (mean TPM = 3.6 vs mean normal TPM = 39.12), and in unaffected cancers (mean TPM = 21.85, Figs. S2a,b). However, we did not observe a systematic decrease in expression across all gene-patient combinations (Fig. S2b), likely due to other factors (copy number alterations, phenotypic plasticity, undetected SCAAs) influencing expression.

**Figure 2.**
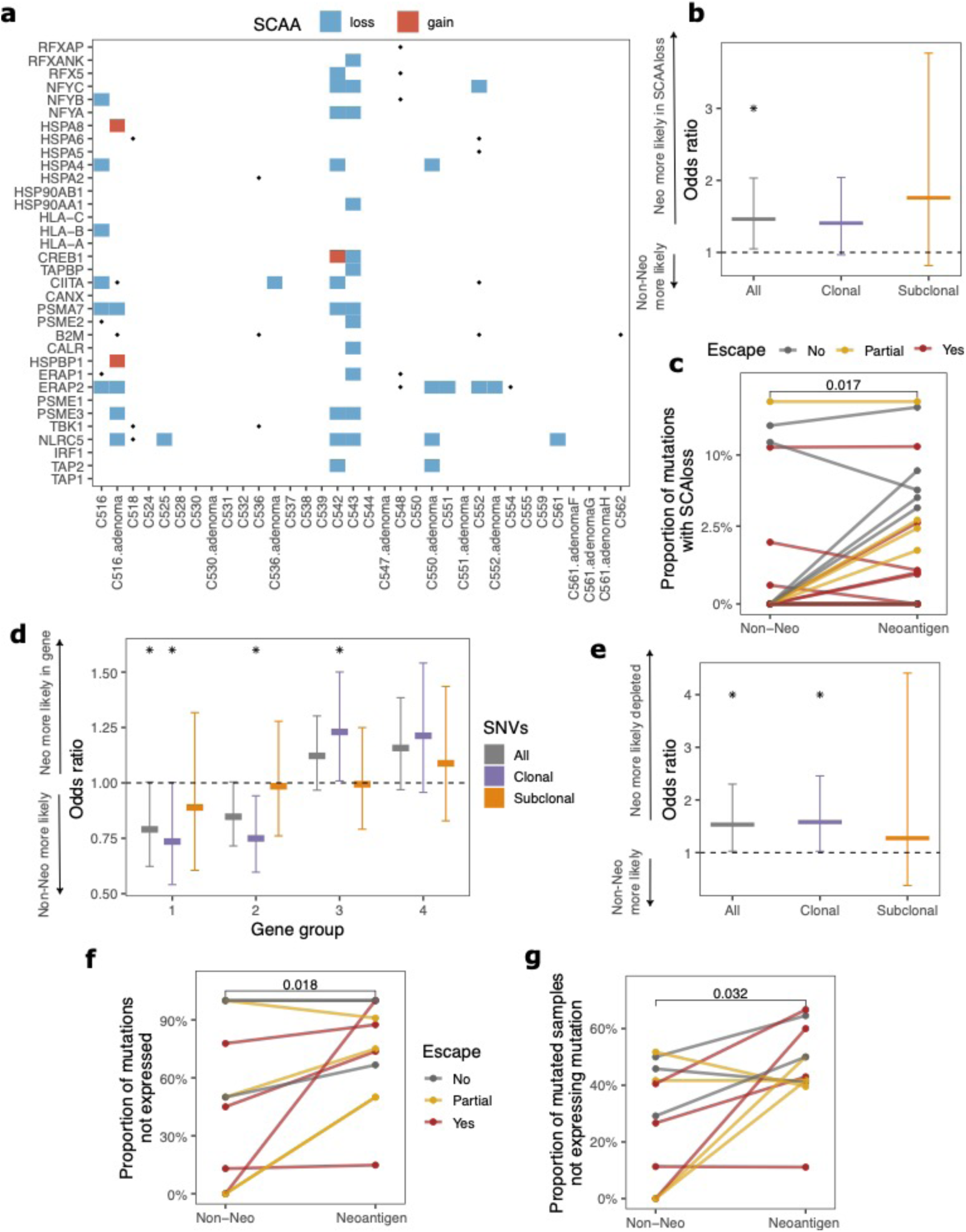
SCAAs and transcriptional regulation decrease neoantigen presentation. (a) Heatmap of SCAAs in antigen presenting genes. Blue and red rectangles show SCAA losses and gains in gene-associated promoter regions, with dots indicating somatic mutations. (b) Odds ratio of a neoantigen vs non-antigenic mutation being located in a gene affected by promoter SCAAloss. (c) Proportion of mutations identified by WGS that are located downstream of a SCAAloss promoter. (d) Odds ratio of a neoantigen vs non-antigenic mutation being located in genes of gene group 1-4 from ref^21^. All, clonal and subclonal mutations are shown separately in grey, purple and orange. (e) Odds ratio of a neoantigen vs non-antigenic mutation being transcriptionally edited (present in WGS but not expressed in RNAseq). (f) Proportion of mutations found in WGS that are edited in RNA for each FF-WGS cancer with matched RNAseq data. (g) Proportion of samples within a cancer that carry a given (antigenic or non-antigenic mutation) in WGS but not in corresponding RNAseq. In c,f&g, cancers are coloured according to their immune escape status.

In our prior work, we found that gene expression in CRCs shows high plasticity and low heritability^21^. Motivated by these findings, we examined the heritability of expression within APGs specifically (see Methods) using phylogenetic signal analysis^25,26^. Nine cancers had sufficient number of biopsies with matched WGS and RNAseq for this analysis. The strongest phylogenetic signal (correlation between expression and evolutionary distance) was that of HLA-A expression in patient C559, which had a clade of biopsies with subclonal LOH in HLA-A. TAPBP and HSP90AB1 were the only genes recurrently phylogenetic genes although we detected somatic mutations in neither (Figs. S1 & S2c). However, in patient C559 we detected differential patterns of open chromatin between tumour regions B and C&D (region A had no ATAC-sequenced samples) in the regulatory region upstream of HSP90AB1 (Fig. S8d), making it plausible that heritable epigenetic variation is behind expression regulation. The generally low level of phylogenetic signal (signal found only in 17/297 gene-cancer combinations evaluated), combined with high intra-tumour heterogeneity (expression differences on the magnitude of median expression) confirms that the expression of antigen presenting genes is plastic, similar to the majority of genes.

### Immuno-editing through SCAAs and transcriptional regulation

We examined whether epigenome reorganisation was responsible for preferentially silencing neoantigens, leading to epigenetic immuno-editing. We collated the chromatin accessibility of all genetic loci where a protein-changing mutation was detected. Neoantigens were enriched in genes where SCAAs tended to close chromatin (Fig. 2b, OR_(neoantigen & SCAAloss)_=1.46[1.05-2.03]), and this observation was true on the individual cancer level as well (Fig. 2c, p=0.017). We also collated all SCAAlosses and confirmed that significantly more of these were associated with neoantigens than non-antigenic SNVs (Fig. S3a, p=0.0085).

Next, we examined transcriptional regulation as an additional source of neoantigen expression modulation. In our EPICC cohort, we had previously clustered genes into four gene groups according to each gene’s expression level across and variability within cancers^21^, where group 1 was the most highly and uniformly expressed, and group 4 consisted of lowly expressed genes. For each group, we compared the proportion of neoantigens to non-antigenic protein-changing mutations (neoantigen ratio) falling within these genes. Clonal neoantigen ratios were significantly lower in gene groups 1 and 2 (Fig. S3b), while subclonal neoantigen ratios were similar for all gene groups (Fig. S3c). Groups 1&2 showed a significant depletion of clonal SNV neoantigens (OR_(neoantigen & in group 1)_ =0.73[0.54-1.0] and OR_(neoantigen & in group 2)_=0.75[0.6-0.94], Fig. 2d). To confirm that this depletion was related to expression patterns, we identified consistently expressed genes in our cohort using the definition of Rosenthal et al.^18^ (>=1TPM in >95% of the measured tumour samples). Indeed, SNV neoantigens were significant depleted within consistently expressed genes, but subclonal mutations showed no such difference (Fig. S3d). Frameshift (FS) neoantigens, especially clonal FS mutations, showed an even stronger depletion in gene groups 1 and 2 and in consistently expressed genes (Fig. S3e-h). Thus, clonal neoantigens are typically found in genes with low and/or variable expression.

We also examined allele-specific expression of neoantigens. Clonal neoantigens were significantly less likely to be expressed (OR_(neoantigen & not expressed)_=1.53[1.03-2.3]) than non-antigenic clonal SNVs, while subclonal mutations showed no significant difference (Fig. 2e), and this was also true on the level of individual cancers (Fig. 2f). Further, neoantigen silencing was more wide-spread within a cancer (observed in more biopsies from that cancer) than silencing of non-antigenic SNVs (Fig. 2g). Overall, around 50% of neoantigens (where there were sufficient RNA reads) were found to be transcriptionally immuno-edited, with the exception of C516, a highly escaped MMRd cancer (Fig. S1).

Collectively, we observed that (clonal) neoantigens that survive to be detectable in cancer DNA are enriched in genes with low expression and/or in regions of closed chromatin, and are further depleted through allele-specific expression modulation. All these processes of post-genetic immuno-editing are likely to have an underlying somatic epigenetic mechanism.

### Immune exclusion and suppression across spatially diverse tumour regions

We next examined how the structure and composition of the tumour microenvironment (TME) contributed to immune evasion. Quantitative analysis of cell counts in our matched CyCIF data showed that overall tumours were significantly depleted of immune cells (quantified by the number of immune cells per epithelial cell, Fig. 1c). The fraction of CD8+ cytotoxic T-lymphocytes (CTLs) was significantly lower in superficial tumour and in the invasive margin than in adjacent normal mucosa (Fig. 3a, p=3×10^−5^ and p=3×10^−3^, respectively) and CTLs were more distant from epithelial cells in tumour-containing regions than in normal mucosa (Fig. 3b-c, p<10^−16^). CTL-tumour cell distance was highest in superficial tumour regions. Further, CTLA4-expressing FOXP3+ regulatory T-cells, associated with an immune-suppressive function, were enriched in all tumour regions compared to normal mucosa (Fig. 3d; p=0.012, p=0.01 and p=6×10^−3^ for superficial tumour, invasive margin, and node), consistent with previous reports^28^.

**Figure 3.**
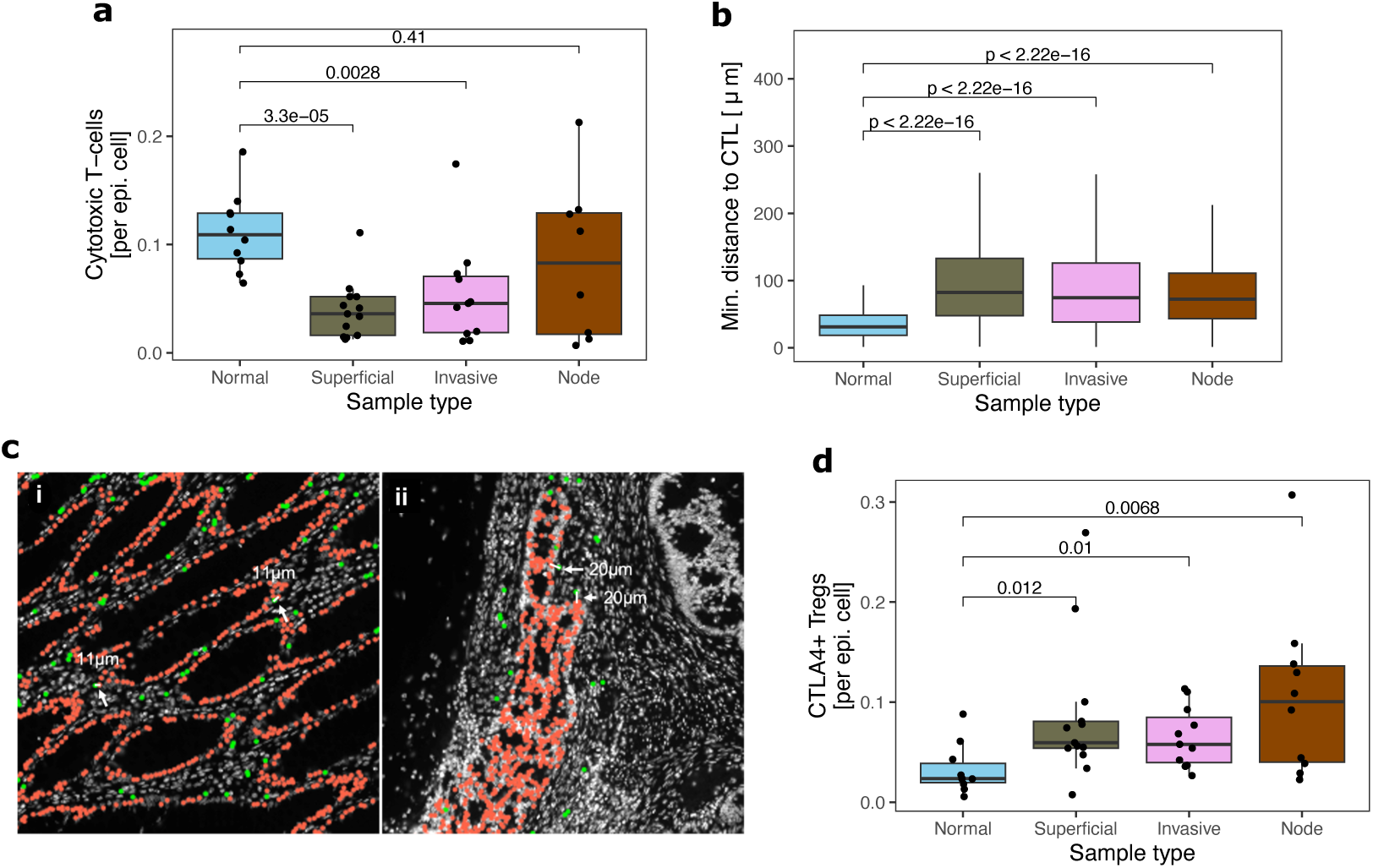
Immune exclusion and clonal immune escape in CRCs. (a,d) Number of cytotoxic T-cells (a) and CTLA4+ Tregs (d) normalised to epithelial cells in each CyCIF image, grouped by sample type. (b) Distance between epithelial cells and the closest cytotoxic T-cell in different sample regions (n=66809, 122280, 180485, 124762 in normal, superficial, invasive and node, respectively). (c) Representative CyCIF images showing epithelial cells (in red) and cytotoxic T-cells (in green) in normal mucosa (i) and in superficial tumour (ii).

We then examined serial haematoxylin and eosin (H&E) stained slides with a deep learning-based cell classifier^29^ that identified tumour cells, lymphocytes and a number of other cell types. We confirmed that the fraction of lymphocytes was significantly reduced in superficial tumour samples compared to both adjacent and distant normal mucosa (Fig. S4, p=1.5×10^−6^, p=1.6×10^−3^, respectively). However, the total lymphocyte count in the invasive margin and lymph node deposits was similar to that observed in normal samples, indicating only CTLs, but not other lymphocyte types, were depleted at invasion.

### Prevalence and consequence of genetic immune escape

We examined the evolution of genetic immune escape alterations. Clonal immune escape mutations were detected in 8/29 CRCs (Figs. 1b & 4a,b, Fig. S1) with three cancers (C518, C524, C548, Figs. 4c,d & S1) carrying multiple HLA alterations on independent branches of the phylogenetic tree. This parallel emergence suggests strong selection for immune escape^30^. Minor subclones with immune escape alterations (alteration present in less than 25% of samples from a tumour) were detected in a further 4 cases (C537, C543, C547, C559; Figs. 1b & S1). Two HLA mutations (in C524 and in C537) were also confirmed in the invasive margin of matched FFPE-PS samples.

**Figure 4.**
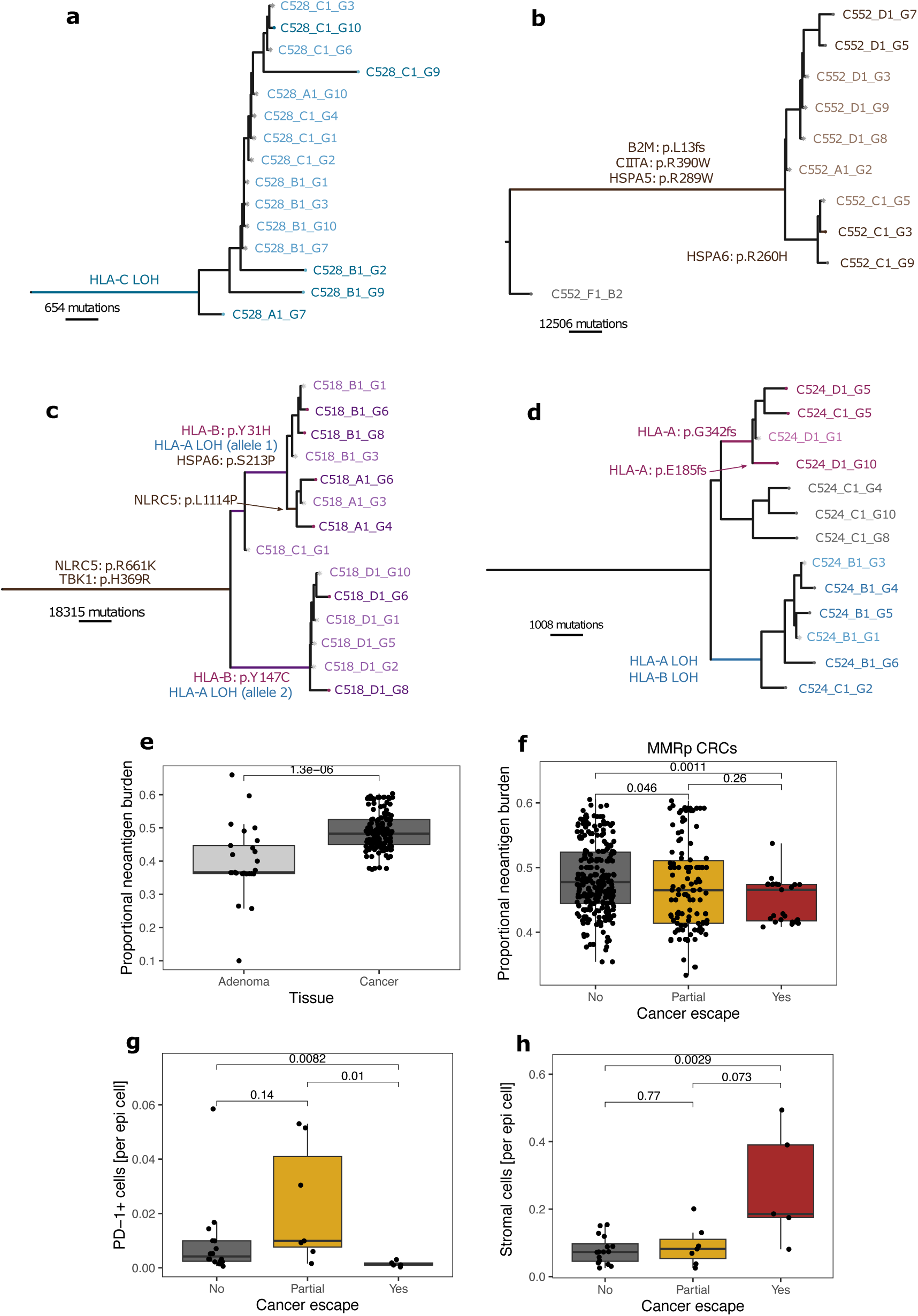
Association between timing of immune escape, -editing and exclusion. (a-d) Phylogenetic tree reconstructed using FF-WGS data of patient C528 (a, MMRp), C552 (b, MMRd), C518 (c, MMRd) and C524 (d, MMRp). Immune escape alterations are shown by arrows or over the branch they occurred in, with colours representing mutations as in Fig. 1b. Light-coloured samples were sequenced using low-pass WGS and genotyped using deep-sequenced samples. (e) Proportional burden of CRA and CRC biopsies. (f) Proportional burden values for MMRp biopsies, according to the immune escape status of each cancer. (g-h) Number of PD-1+ (g) and stromal (h) cells per epithelial cell in FFPE-PS samples, according to cancer immune escape status.

We found that alterations with the highest predicted impact on immune evasion (based on prior studies) were mostly shared across the whole tumour or multiple spatially distinct regions, meaning that they were early evolutionary events. For example, beta-2-microglobulin (B2M) mutations were all clonal (n=4), predicted to highly impact transcription/function through start-loss or frameshift (Fig. 4b & Fig. S1). Matched RNAseq revealed significantly decreased B2M expression in these cancers confirming the impact of these mutations (Fig. S5a). Similarly, we found mutations in NLRC5 and RFXAP – essential factors in the MHC class-I enhanceosome^31^ that reduced expression of class-I MHC genes (HLA-A, -B, and -C, Fig. S5b,c) – were also shared across multiple regions. HLA LOH, a common immune escape mechanism shown to be more impactful than HLA mutations^32,33^, was also clonal or near-clonal in 4/5 cancers with HLA-LOH (Fig. 4a & Fig. S1). In contrast, likely low-impact (SNV) mutations in antigen presentation showed the highest variability within tumours: we detected 8 HLA SNVs and 5 SNVs in other APGs that were unique to a single gland or tumour region. We suggest that these alterations likely confer only limited disruption to neoantigen presentation and are weakly selected.

To examine the timing of immune evasion during initial cancer formation, we evaluated neoantigens and immune escape variants in 25 glands from eight colorectal adenomas (CRAs), including an advanced cancer-adjacent MMRd adenoma from patient C516. No adenoma glands carried immune escape mutations (Fig. S1), with the exception of the advanced CRA in C516 (Fig. S1a). Further, CRAs had significantly lower proportional neoantigen burden than CRCs (Fig. 4e), indicating that immune surveillance is more active and effective in CRAs than in CRCs. Therefore, it is likely that immune escape occurs at the outset of, or early during, CRC outgrowth.

We measured how the presence and extent of immune escape correlated with neoantigen burden and immune selection. In MMRp CRCs, the presence of genetic immune escape was associated with a lower overall (clonal & subclonal combined) proportional neoantigen burden (Fig. 4f), possibly because these cancers had been stringently immuno-edited prior to immune escape. Noteworthy was that C524 and C548, cases with parallel evolution of HLA alterations, had biopsies with immune dN/dS<1 (strigent immunoediting) suggesting past stringent immune surveillance that led to selection for these HLA alterations (Fig. 1b).

In the matched CyCIF data, PD1+ cells and VISTA+ cells, that are known to mediate T-cell exhaustion^34^, were enriched in partially escaped cancers (Fig. 4g & Fig. S5d), indicating a TME with impaired immune elimination ability. Fibroblasts (stromal cells) were more abundant in escaped cancers, while CTLs and Tregs showed no difference by escape status (Fig. 4h & Fig. S5e-f). Partially escaped CRCs also had an intermediate overall proportional neoantigen burden, with immune dN/dS values below 1 indicating immunoediting (Fig. S5g). These findings suggest that partially escaped cancers were subject to strong selection during early development, prior to gaining subclonal escape.

Both proportional burden and immune dN/dS detect selection through the absence/decreased number of antigenic mutations, and therefore detects only selection that leads to mutation elimination. For contrast, we assessed neoantigen variant allele frequency (neoantigen-VAF) that can detect antigenic clones which are neoantigen depleted, but not eliminated, indicative of weak or ongoing selection^8^. While we did not observe consistent neoantigen-VAF depletion across (superficial) samples (Fig. S6a-b), significant depletion was observed when analysis was restricted to escaped MMRp cases (Fig. S6c, KS-test p=0.012). Similarly, neoantigens were as likely to be shared between samples from a tumour as non-antigenic mutations (Fig. S6d-f). These data are indicative of relatively uniform immune selective pressures across large tumour regions.

### Intra-tumour heterogeneity in immunoediting

We sought to explore intra-tumour heterogeneity in genetic immunoediting using our spatially resolved gland-level samples. We examined the correlation between histological region and neoantigen burden in comparison to the presence/absence of immune escape and other gland-to-gland tumour-intrinsic difference. The genetic similarity of samples correlated with the physical distance between samples (Fig S7a,b). Multivariable regression (Methods) showed that the vast majority of neoantigen burden variation was explained by patient specific effects and progression status (CRA vs CRC), with negligible contribution from other variables (Fig. 5a & Fig. S7c). In FFPE-PS samples, patient-specific effects also dominated over the histological location of the sample (superficial, invasive edge or from a nodal lesion) (Fig. 5b & Fig. S7d). Using the immune-dNdS measure in place of neoantigen burden gave analogous results (Fig S7e-f). This was also true when clonal and subclonal mutations were considered separately, and when tumours were stratified by immune escape status (Fig S7g-i).

**Figure 5.**
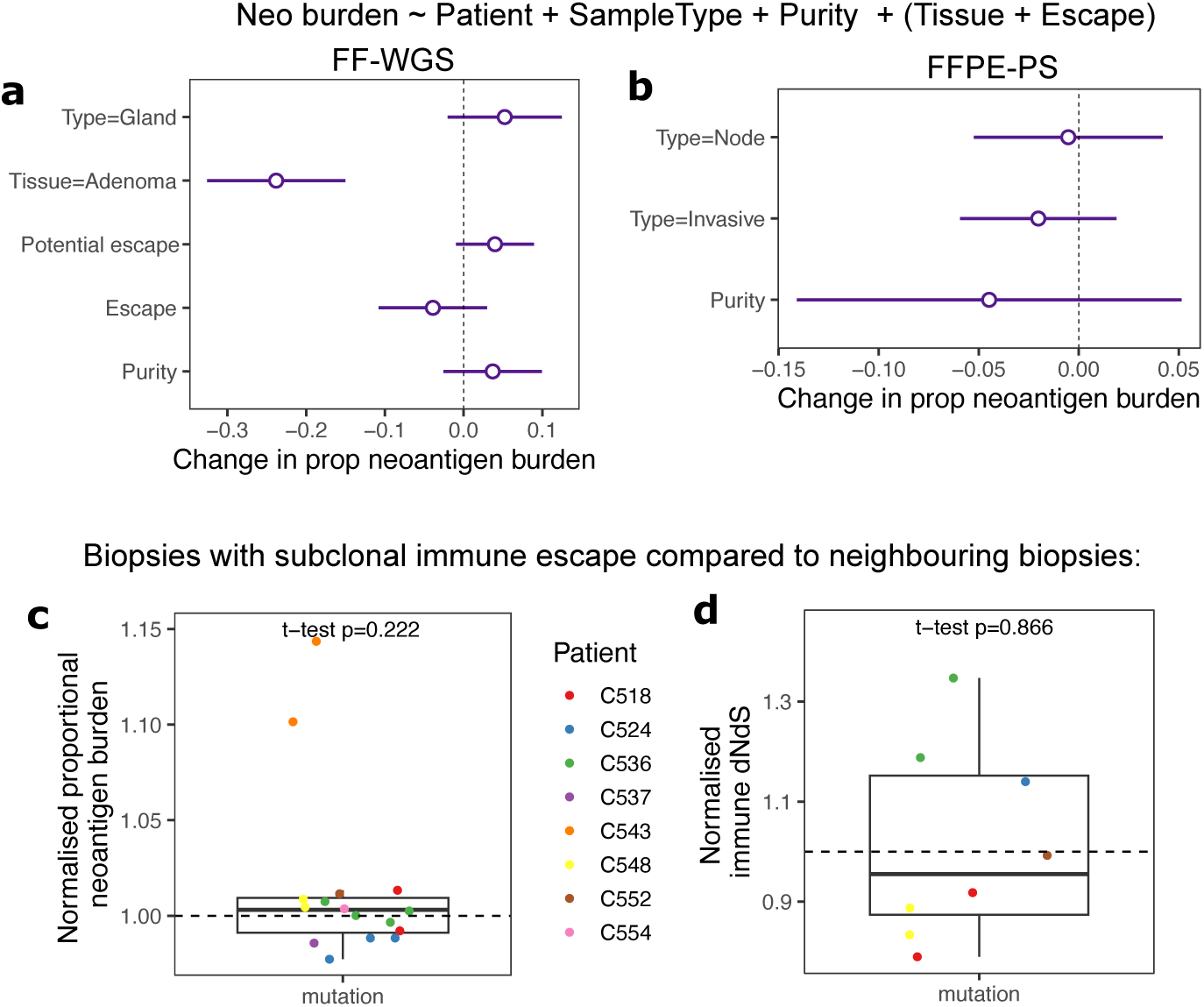
Intra-tumour differences in immune selection. (a-b) Forest plots depicting the output of a multivariable regression (Methods) investigating the association of proportional neoantigen burden with other sample characteristics in FF-WGS (a) and FFPE-PS (b) samples. Circles denote the estimated coefficients, with whiskers showing 95% confidence intervals. (c-d) Proportional neoantigen burden (c) and immune dNdS (d) in FF-WGS biopsies with subclonal immune escape. Values have been normalised by the average of all deep-sequenced biopsies phylogenetically close to the escaped biopsy (see Fig. S1). Individual cancers are depicted in different colours. T-tests against the null hypothesis of mean=1 are shown on top of each panel.

We analysed the impact of subclonal immune escape on by comparing neoantigen burdens between escaped and non-escaped regions in cancers with subclonal immune escape mutations (C518, C524, C536, C537, C543, C548, C552, C554, Fig. S1). Subclonal escape was not systematically associated with higher normalised neoantigen burden or immune dNdS values, with the exception of HLA-mutated biopsies in C543 that did carry a higher antigenic burden (Fig. 5c-d). These data confirm that subclonal escape had at most a modest effect on the intensity of immunoediting.

### Localised tumour-immune interactions at the invasive margin

We focused on immunoediting and TME structure at the invasive margin and lymph node deposits. We divided lymphocytes into tumour-infiltrating, -adjacent or -distant^35^, and observed that invasive margin samples, but not node deposits, were characterised by a higher ratio of infiltrating lymphocytes compared to the superficial tumour (Fig. 6a). Using our CyCIF data, we observed that the proportion of PD-L1+ tumour cells (that are likely repressive of T-cell activity) was significantly higher at the invasive margin than in superficial tumour (Fig. 6b). These results imply heightened immune surveillance at the invasive margin that is then reduced in node deposits.

**Figure 6.**
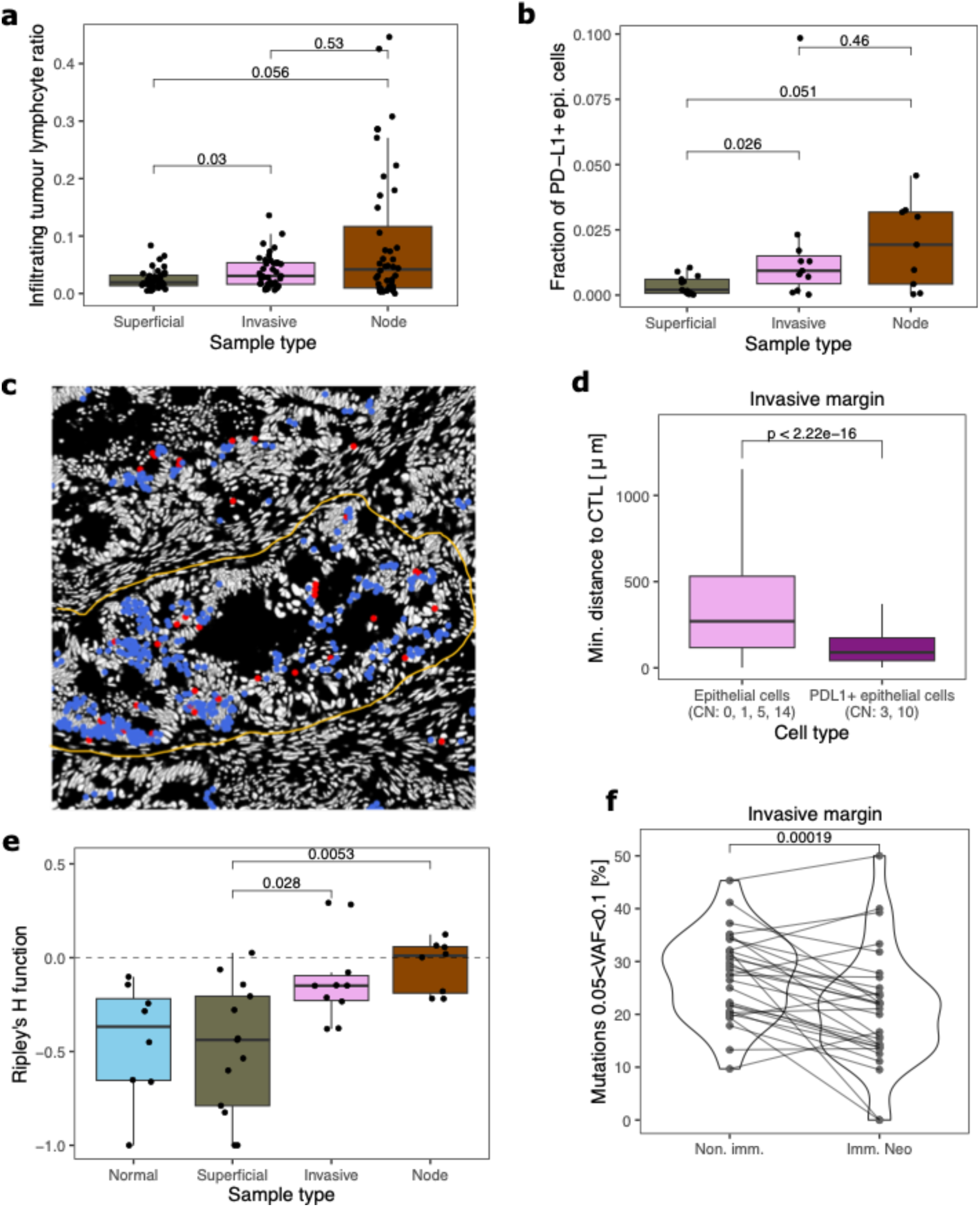
Increased immune interaction and immuno-editing in the invasive margin. (a) Infiltrating tumour lymphocyte ratio quantified using H&E stained images, according to the type of tumour-associated region of interest. (b) The fraction of PDL1+ epithelial cells in CyCIF images across different sample types. (c) Representative CyCIF image of a tumour gland from the invasive margin, showing PD-L1+ epithelial cells in blue and cytotoxic T-cells in red. The outline of the gland is shown by a yellow line. (d) Distance of epithelial cells to the closest cytotoxic T-cell, for non-PDL1-expressing epithelial cells in non-PD-L1-CNs and for PD-L1+ epithelial cells in PD-L1-associated CNs (n=158547, 4459 for non-PD-L1 and PD-L1+ cells, respectively). (e) Ripley’s H function measuring intermixing between PD-L1+ and PD-1+ cells across different cancer regions. Values around zero (dashed line) indicate uniform mixing. (f) The proportion of all high-immunogenicity neoantigens and of all non-immunogenic mutations falling within the range 0.05<VAF<0.1 in invasive margin samples. Each line indicates paired values derived from the same biopsy.

We sought additional evidence of heightened immune surveillance at the invasive edge. We derived cellular neighbourhoods (CNs) from CyCIF data by considering motifs of an index cell and its ten nearest neighbours^36^, identifying 15 distinct CNs (Fig. S8a). The fraction of cells belonging to CN3 (highly enriched for PD-L1+ cells, including PD-L1+ epithelial cells) was significantly higher in invasive margin and node deposits than in superficial tumour (Fig. S8b). Of note, CN10 also had PD-L1+ cells and other immunosuppressive cells, including PD-1+ and CTLA4+ cells, but this neighbourhood was present at similar proportions in all tumour-associated regions (Fig. S8c).

PD-L1+ tumour cells were dispersed within PD-L1+ neighbourhoods, even within single tumour glands (Fig. 6c), suggesting high plasticity in expression of this molecule. To investigate the functional drive behind PD-L1+ expression, we examined co-localisation of CTLs and PD-L1+ cells by computing the distance between each PD-L1+ epithelial cell and its closest CTL. We found that PD-L1+ epithelial cells were significantly closer to CTLs than PDL1-negative epithelial cells (Fig. S8d), with the closest PD-L1+ CTL relationship observed in the invasive margin (Fig. 6d & S8e). Moreover, PD-L1+ cells were mixed uniformly with (cognate) PD-1+ cells in the invasive margin and node, while they often showed exclusion in superficial tumour and normal tissue (Fig. 6e). These observations suggest that CTL surveillance is heightened at the invasive edge and PD-L1 expression may arise as an adaptive response to it.

In matched genomic data, we explored immune selection signal within the invasive margin samples specifically. We observed that these samples contained candidate highly immunogenic neoantigens at lower VAFs than non-immunogenic mutations (see Methods for definition, Fig. S9a, KS-test p=6×10^−4^). Depletion of neoantigens was most prevalent within small subclones (0.05 < VAF < 0.1, Fig. 6f, p=2×10^−4^). On the other hand, no neoantigen VAF depletion was observed in superficial tumour or node deposits (Fig. S9b-c), or for low-antigenicity neoantigens (Fig. S9d), in agreement with our earlier finding that sample location was not associated with systematic differences neoantigen elimination. Therefore, it appears that enhanced immuno-editing occurs as a variable ongoing process at the point of invasion, primarily affecting very small subclones (~100 cells) of increased immunogenicity.

## DISCUSSION

In this work, we mapped the interplay of epigenetic, genetic microenvironmental processes that collectively establish immune evasion and shape tumour-immune coevolution in colorectal cancer. We have shown that these immune evasion processes are part of the “Big Bang” that forms CRCs^42^, and determine the immunogenicity of the whole cancer.

We identified somatic chromatin accessibility alterations (SCAAs) as a mechanism that contributes to the down-regulation of neoantigen-carrying genes and genes associated with antigen presentation. Further, immunogenic alterations, especially clonal neoantigens, are preferentially depleted in transcription through (so far uncharacterised) epigenetic mechanisms. Transcriptional down-regulation is mainly limited to clonally shared neoantigens, reiterating that immune selection is primarily directed at clonal antigens^37,38^. Previous work suggests that low or intermittent expression of antigens can lead to desensitisation of T-cells against these antigens^39^, potentially explaining why transcriptional regulation is more prevalent than genetic depletion in our data. Our results align with previous observations that epigenetic and environmental factors play a substantial role in determining cancer prognosis and response to immunotherapy^18,40,41^. Our finding suggests that epigenetic control could play a key role in both MMRp and MMRd cancers in facilitating resistance to immune surveillance. Consequently, mapping the epigenome could identify patients with lower/higher potential for immunotherapy response. Feasibly, epigenome modifying drugs could have a profound impact on neoantigen presentation and potentially synergise with immunotherapeutic drugs.

Whilst there is heterogeneity in the neoantigen burden and composition of the immune microenvironment, we observed that the features that are inherent to the tumour as a whole are most important for determining immunogenicity. We found that the acquisition of immune tolerance is essential for the adenoma-to-carcinoma transition, and in cancers typically multiple mechanisms of immune evasion occur in tandem. Microenvironmental remodelling is prevalent throughout all tumour-associated regions, coupled with high-impact immune escape genetic alterations and epigenetic processes to ensure reduced neoantigen presentation and immune activation. We observed decreased neoantigen burden in MMRp cancers with immune escape (as opposed to non-escaped MMRp cancers), suggesting that immuno-editing precedes the acquisition of immune escape alterations, and is a key determinant of the clonal neoantigen burden of these cancers. On the other hand, immune-tumour co-evolution past transformation, and subclonal changes to the immunogenomic landscape of the tumour have only a weak influence on neoantigen composition. Collectively, these data suggest CRC are no longer engaged in an “all-out war” with the immune system after initial expansion.

We found that ongoing “battles” of cancer-immune interaction are limited to small localised subclones along the invasive margin of MMRp CRCs, where we presume that there is a large sudden change in immune microenvironmental composition and hence more pronounced immunoediting. These subclones are characterised by an intricate interplay of higher CTL infiltration, depletion of subclonal neoantigens and PD-L1+ expression, painting the picture of an ongoing skirmish between active immune and cancer cells. However, we did not observe any variation associated with subclonal immune alterations within the superficial tumour, confirming that the tumour bulk is not shaped by such ongoing battles.

Overall, our work indicates the acquisition of immune escape and/or an immune excluded microenvironment is part of the “Big Bang” necessary for the adenoma-to-carcinoma transition in CRCs. We observe that epigenetic, transcriptomic and microenvironmental mechanisms add a layer of regulation to the immunogenomic profile of CRC cells, allowing further adaptation through both hereditary and plastic means. Our findings suggest evaluating and targeting of epigenetic machinery and TME reorganisation as a promising new way to stratify patients and enhance the efficacy of immunotherapy.

## ACKNOWLEDGEMENTS

This work was supported by the Wellcome Trust (202778/Z/16/Z to T.A.G.; 202778/B/16/Z to A.S.; 105104/Z/14/Z to the Centre for Evolution and Cancer, Institute of Cancer Research) and Cancer Research UK (A19771 and DRCNPG-May21_100001 to T.A.G. partially supporting E.L.; A22909 to A.S.; Clinical Research Training Fellowship supporting V.G., Accelerator Award A26815 supporting A.S.). E.L. is supported by a start-up grant from the Chalmers Area of Advance Health Engineering. V.G. received support through a Clinical Lecturer Grant from the Jean Shanks/ Pathological Society (JSPS/CLG/1022/01). This work was also supported in part by NIH grants U54 CA217376, U2C CA233254, and R01 CA140657. The findings, opinions and recommendations expressed here are those of the authors and not necessarily those of the universities where the research was performed or the National Institutes of Health.

## AUTHOR CONTRIBUTIONS

A.S. and T.A.G. conceived the study and A.S., T.A.G. E.L. and V.G. designed research. C.C.M., A.S and T.A.G. acquired funding for the project. E.L., V.G. and T.A.G. wrote the original manuscript with contribution from A.M.B. M.M., I.S., C.K., M. Mitchison, M.J., M. R.-J. and J.B. contributed to sample collection and data generation. V.G. performed FFPE-PS sequencing and CyCIF analysis with support from A.M.B. E.L. performed multi-omic FF-WGS analysis. V.G. and E.L. performed FFPE-PS bioinformatic analysis. L.Z., J.H., T.H., C.L., G.D.C. and G.L contributed to bioinformatic analyses. N.T., O.S. and L.C. contributed to imagine analysis. All authors reviewed and approved the manuscript.

## METHODS

### Code and data availability

Scripts required to reproduce data processing and the figures of the manuscript is available at https://github.com/elakatos/EPICC_immune_analysis. Processed data used in the figures and to derive summary tables are available at Mendeley: https://doi.org/10.17632/cjfmmc95dm.1. Raw sequencing reads of FFPE-PS samples will be deposited on the European Genome-Phenome Archive. For raw sequencing data of FF-WGS samples, see refs^15,21^.

### FF-WGS sample collection and sequencing

Sample collection and processing is detailed in our previous works describing the evolutionary predictions in colorectal cancer (EPICC) cohort in refs^15,21^.

Briefly, primary tumour tissue and matched blood samples were prospectively collected and four regions from each primary cancer were sampled by punch biopsy or scalpel dissection, further divided and slow-frozen to −80 °C. Individual glands were isolated under microscope, and when these were not available, epithelial minibulks comprised of 10-20 crypts/glands were collected. RNA and DNA were extracted, and WGS and RNA libraries were prepared using the NEBNext Ultra II FS kit and Illumina TruSeq RNA Exome kit, respectively. Sequence libraries were multiplexed and sequenced on NovaSeq (Illumina).

### FFPE-PS sample collection and sequencing

11 stage-III microsatellite-stable cancers which had lymph node metastases were identified from the EPICC cohort, 9 of which have been included in the FF-WGS cohort. FFPE sections were cut in the following order: H&E-1 (5μm), 5x 8μm for laser capture microdissection, H&E-2 (5μm), 8x 5μm for cyclic immunofluorescence, H&E-3 (5μm). The H&E slides from each FFPE block were digitised using the NanoZoomer S210 or S60 (Hamamatsu, Welwyn Garden City, UK). Images were reviewed using NDPViewer software.

Regions of interest (ROIs) were identified as single glands or clusters of small adjacent glands (micro-biopsies) from superficial, invasive margin or lymph node deposits. Superficial regions were defined as cancer regions adjacent to or contiguous with normal mucosa. The tumour-normal interface was identified where possible, and the invasive margin was defined as the region within 500μm either side of the tumour-normal interface (with an overall extent of ~1mm).

#### Panel sequencing

ROIs were micro-dissected using PALM MicroBeam Laser Microdissection (Zeiss). DNA was extracted using the High Pure FFPET DNA Isolation Kit (Roche). Extracted FFPE DNA was repaired using the NEBNext FFPE DNA Repair Mix. Post-repair, whole genome libraries were prepared using the NEBNext Ultra II FS DNA Library Prep Kit for Illumina with unique molecular identifier (UMI) adaptors ligated onto DNA molecules.

Panel sequencing on FFPE samples was carried out using a custom targeted panel designed by Dr Zapata and manufactured by Twist BioSciences, focusing on regions encoding the immunopeptidome and antigen-presenting related genes. The immunopeptidome was defined as the set of the human 9-mers that strongly bind (rank <0.5 according to netMHCpan 4.0) one of the top 70 HLA alleles, was confirmed T-cell positive in IEDB and were derived from a gene with mean expression >1FPKM pan-cancer. The final list of these immunopeptidome loci can be obtained as a bed file at: https://github.com/luisgls/SOPRANO/blob/master/immunopeptidomes/human/allhlaBinders_exprmean1.IEDBpeps.unique.bed.

The Twist Target Enrichment Protocol was used to hybridise probes from the this custom panel with prepared libraries. Hybridised targets were isolated and amplified with PCR. Paired-end 50bp runs were performed on Novaseq S1 (Illumina).

### Processing of sequencing data

Alignment of fastq reads produced for FF-WGS data, copy number analysis and SNV detection are detailed in ref^15^.

For FFPE-PS samples, three sets of fastq reads were aligned to human reference genome build hg38 to generate an unmapped BAM using FastqToBam (Fgbio 1.3.0). Fastq files were then created from this unmapped bam (SamToFastq, Picard version 2.20.3) and aligned to human reference genome build GRCh38/hg38 with Burrows-Wheeler Aligner package BWA-MEM version 0.7.17. Alignment data from the outputted bam from this step was merged with data in the previously generated unmapped bam (MergeBamAlignment^170^, Picard version 2.20.3). The merged bam file was then used as input for GroupReadsByUMI (Fgbio 1.3.0). Reads were initially grouped by template and then reads which had the same end positions were sub-grouped by UMI sequence. “Adjacency” was used as the strategy for grouping so that errors were allowed between UMIs, but only when there was a count gradient. The allowable number of edits/ changes between UMIs was set to 1 (default). Minimum mapping quality for mapped reads was set to 30 (default).

Consensus sequences were called from reads with the same unique molecular tag (CallMolecularConsensusReads, Fgbio 1.3.0) and filtered using FilterConsensusReads (Fgbio 1.3.0) to exclude consensus sequences with fewer than 2 contributing reads, mask consensus bases with quality less than 30, accept a maximum raw-read error rate across the entire consensus read of 0.05, accept a maximum error rate for a single consensus base of 0.1 and accept a maximum fraction of 0.2 for no-calls in the read after filtering.

The filtered reads were then ultimately aligned to human reference genome build GRCh38/hg38, and variant calling was performed using Mutect2 (GATK) with a bed file specifying regions of the genome covered by the targeted panel. Normal samples from adjacent muscle were used as matched normal for variant calling. The resulting vcf files were merged for each patient and passed to Platypus for multi-region variant calling.

Variant calls were filtered to retain mutations within certain filtering criteria and adequate support for the variant, as described in ref^15^. The same filters were used for both FF-WGS and FFPE-PS samples, except for requiring >=5 and >=8 reads covering each site in FF-WGS /FFPE-PS samples respectively.

FF-WGS samples sequenced at shallow depth were genotyped and fitted on phylogenetic trees using the method described in ref^15^.

### HLA haplotyping

HLA-A, -B and -C haplotyping was performed using polysolver^43^ on FF-WGS samples, by running shell_call_hla_type with default settings. As ethnicity information was not available, we used “Unknown” for all samples. To increase coverage over the HLA gene regions, we used merged bam files created from all sequencing bam files from a given patient. For validation, we also performed haplotyping on merged bams formed of normal (blood or normal colon tissue) samples and compared the predicted haplotypes. The predicted haplotypes had a high concordance, with haplotypes predicted using all samples providing one more heterozygous haplotype than normal-only haplotypes in 6/30 cases. Based on the average homozygosity across colorectal cancers (as seen in TCGA CRC samples^8^), we accepted the more heterozygous set of alleles predicted to define their set of HLA alleles that was inputted into neoantigen prediction and HLA alteration prediction. For all HLA alterations, we confirmed that the alterations are called independent of which haplotyping calls were used.

For FFPE-PS samples we used the calls derived from matched FF-WGS samples. For the two patients where this was not available, we performed haplotyping on adjacent normal mucosal samples following the same steps as described above.

### Immune escape prediction

#### Mutations in HLA

Somatic mutations in the HLA locus were predicted using polysolver^43^. The mutation detection script of polysolver (shell_call_hla_mutations_from_type) was run on matched tumour-normal pairs to call tumour-specific alterations in HLA-aligned sequencing reads (using the haplotypes previously predicted) using MuTect (v1.16). In addition, Strelka2^44^ (v2.9.10) was independently run to detect short insertions and deletions in HLA-aligned reads as this version offers increased sensitivity over polysolver’s default caller. Both single nucleotide mutations and frameshift alterations passing quality control were annotated by polysolver’s built-in annotation script, shell_annotate_hla_mutations. Based on this annotation, a mutation in the HLA locus was called if a mutation passed all quality filters and introduced either a missense/nonsense change or was located at a splice site. In addition, we also identified second-tier mutations in unfiltered MuTect files that were detected in insufficient reads to pass quality control, but the exact same nucleotide change was clearly detected in another biopsy of the same tumour. This way we could identify mutations that would otherwise be masked by lower tumour purity and sequencing depth.

#### Loss-of-heterozygosity in HLA

LOH at the HLA locus was predicted using LOHHLA^32^ and sequenza^45^. First, we evaluated the allele-specific copy number as predicted by sequenza at the HLA-A, -B, -C loci. Samples with a predicted minor allele copy number of 0 (e.g. 2:0, 3:0) were labelled as candidate LOH. Then, we ran LOHHLA with the polysolver-generated haplotype files and matched tumour-normal pairs as input. A type-I allele of a patient was annotated as “allelic imbalance” (AI) if the p-value testing the difference in evidence for the two alleles was lower than 0.01. Alleles with AI were labelled as LOH if the following criteria held: (i) the predicted copy number of the lost allele was below 0.5 with confidence interval strictly below 0.7; (ii) the copy number of the kept allele was above 0.75; (iii) the number of mismatched sites between alleles was above 10. HLA LOH was identified by both methods independently in 18 deep-sequenced samples. HLA LOHs called by only one of the methods were manually inspected. Of LOHHLA-exclusive calls, 2 were found to be false positives, and 4 were identified as allelic imbalances (AI) with a minor allele copy number of 1. Of sequenza-exclusive calls, 1 region showed clear LOH and got classified as HLA LOH; 6 samples showed similar CN pattern, but were missed by LOHHLA due to low purity – as adjacent low-pass sequenced samples showed CN=1 in the HLA region, we also classified these as HLA LOH. In addition, 25 regions had allelic imbalance detected by LOHHLA that were also confirmed in sequenza to have a CN state of 2:1 or 3:1.

#### Mutations in antigen presenting genes

We assembled a list of genes involved in class-I type MHC presentation, by using the KEGG pathway ‘antigen processing and presentation’ and MHC I pathway specifically. The following genes were considered: *TAP1, TAP2, IRF1, NLRC5, TBK1, PSME3, PSME1, ERAP2, ERAP1, HSPBP1, CALR, B2M, PSME2, PSMA7, CANX, CIITA, TAPBP, CREB1, HLA-A, HLA-B, HLA-C, HSP90AA1, HSP90AB1, HSPA2, HSPA4, HSPA5, HSPA6, HSPA8, IFNG, NFYA, NFYB, NFYC, RFX5, RFXANK, RFXAP*. Then, we evaluated the expression of each gene within our cohort and filtered out genes that were not clearly expressed (>=10TPM) in at least 5% of samples.

Then, we called evaluated the mutations called in these genes following. Only mutations with at least moderate predicted impact were called. In the overall cohort, we detected mutations in the following genes: NLRC5, TBK1, ERAP2, ERAP1, B2M, PSME2, CIITA, HSPA2, HSPA5, HSPA6, NFYB, RFX5, RFXAP. Mutations with high impact (due to start-loss and frameshift mutations) were detected in B2M and RFXAP.

### Neoantigen prediction and proportional burden computation

We predicted neoantigens from somatic mutation calls and patient-specific HLA haplotypes using NeoPredPipe^24^ for both FF-WGS and FFPE-PS samples. We defined neoantigen burden in a sample as the number of (unique) mutations giving rise to at least one strong-binding (rank <0.5) neoantigen. We evaluated the neoantigen burden arising from both SNVs and frameshift mutations – however, the frameshift neoantigen load of non-MSI cancers was negligible, hence we focused our analysis on SNV neoantigen burden, unless stated otherwise. We also computed the total protein-changing (SNV/frameshift) mutation burden, and used this value to obtain the proportional neoantigen burden for each sample, i.e. what percentage of mutations that has the potential to create a neoantigen does actually lead to strong-binder neoantigens.

#### Defining clonal/subclonal neoantigens

We assigned clonal/subclonal categories to all mutations (independent of neoantigen status) based on their presence/absence in all available deep- or panel-sequenced sample of a given cancer. As the targeted genome region, sequencing strategy and sample types were different, we created separate mutations lists of FF-WGS and FFPE-PS samples. For FF-WGS samples, mutations present in all sequenced cancer biopsies were denoted as clonal and all other mutations (absent in at least 1 but typically more biopsy) as subclonal. For FFPE-PS samples, mutations present in all biopsies were deemed clonal, and mutations absent in at least 2 biopsies were denoted as subclonal.

### Immune dNdS

Immune dN/dS was computed using SOPRANO, with somatic mutation files and personalised immunopeptidome files derived specifically to HLA haplotypes. First, the ratio between dN/dS inside (ON-target dN/dS) and outside the immunopeptidome (OFF-target dN/dS) was computed and corrected for 192-trinucleotide context. Then immune dNdS was computed as the ratio of ON-to-OFF dNdS in order to correct for technical artefacts that could bias dN/dS as computed in OFF-target regions. Samples without any ON- or OFF-target synonymous mutations were excluded from the analysis. In total, immune dNdS estimate was available for 61 FF-WGS and 41 FFPE-PS samples.

To compute immune dNdS separately, we first filtered somatic mutation files to only contain mutations annotated as clonal, then repeated the above procedure on these files.

### VAF distribution of neoantigens/non-antigenic mutations

First, we defined the unadjusted VAF of each mutation as the number of variant reads divided by the number of total reads spanning the locus (Platypus information NV/NR). Then we corrected this value to account for differences in purity: VAF = 1/purity * VAF_unadj_. Purity values for FF-WGS samples were derived from sequenza following manual curation. Purity values for FFPE-PS samples were derived from deep learning classifier applied to the H&E images (see below) overlapping the site resected for sequencing.

Each mutation was annotated according to antigenicity. In FF-WGS samples, we denoted non-binders as non-antigenic mutations and strong-binders as antigens; in FFPE-PS samples we used weak- and non-binders with recognition potential <0.1 as non-antigens and strong-binders with recognition potential >0.1 as antigens, but also explored weak- vs strong-binders (ignoring recognition potential). We only considered SNVs in this analysis. In order to overcome the low numbers of observed mutations, we pulled all mutations from a given set of samples (e.g. samples from escaped MMRp cancers). We then compared the distribution of antigenic and non-antigenic mutations using Kolmogorov-Smirnov test, and visualised both cumulative distributions against the inverse of VAF, looking for the previously established signal of immune selection^8^. In addition, we also evaluated the proportion of antigenic and non-antigenic mutations that were within a certain range of VAF, compared to all antigenic and non-antigenic mutations, respectively. We compared these proportions computed for each sample separately, using a paired Wilcoxon rank sum test.

### SCAA and SCAAloss analysis

We identified somatic chromatin accessibility alterations (SCAAs) by comparing purity- and copy number-corrected ATAC-sequencing peak calls of cancer regions (per cancer) to a pool of normal glands^15^. For immune escape genes, we filtered SCAAs for those located in promoter or enhancer regions associated with a gene from the list detailed in “*Mutations in antigen presenting associated genes*”. Each SCAA was labelled as loss (fold change of cancer compared to normal < −1) or gain (fold change of cancer compared to normal > 1).

To evaluate the observed number of SCAAlosses, we repeated the analysis 200 times with a set of 25-40 randomly chosen genes and computed the number of SCAAlosses and gains. We derived a p-value as the number of random samples we observed a value more extreme than for APGs.

For neoantigen SCAAloss analysis, we filtered SCAA calls to obtain only losses located in promoter regions (within 1000bp of the transcription start site of a gene). Each SCAA loss was annotated by the gene they were proximal to. As the majority of SCA alterations were found to be clonal, we defined SCAAlosses on the cancer level. We then evaluated for each protein-alteration mutation within a cancer whether the gene it is in had an associated SCAAloss or not. For each cancer, we computed the proportion of mutations (both for neoantigens and for non-antigenic mutations) that were classified as downstream of a SCAloss. Similarly, for each SCA loss in a given cancer, we evaluated if it was upstream of a protein-changing mutation and if so. Then within each cancer, we counted the number of SCAlosses upstream of neoantigen/non-antigenic mutations.

### Transcriptional editing analysis

For each SNV detected in FF-WGS samples, we quantified the number of reads in the matched RNA sequencing data supporting the mutation using bam-readcount^27^. For a mutation to be classified as expressed in a sample, we required 3 or more RNA reads overlapping the position to support the variant base. For a mutation to be classified as not expressed (i.e. transcriptionally edited), we required over 10 overlapping reads, with 0 supporting the variant. We chose the threshold >10 so that the probability of misclassifying a mutation present at a true allele frequency of 0.25 or higher was <5%. Mutation-sample pairs that did not qualify for either of these categories were left blank to signify insufficient evidence.

For each cancer, we computed the proportion of mutations that had evidence of transcriptional editing (mutations present in WGS but not expressed in RNAseq) in at least one sample. Similarly, for each mutation in a cancer, we computed the number of samples that expressed that mutation and the number of samples that (confidently) did not express it. We excluded mutations that had sufficient evidence for expressed/not status in less than two samples. We used the median proportion of biopsies with evidence of editing to derive a single value per cancer per mutation type.

### Phylogenetic signal

In order to assess the phylogenetic signal of different immune measures, we followed the procedure detailed in ref^21^. For computing the phylogenetic signal of immune escape gene expressions and mean neoantigen expression, we kept only leaves of each tree that had matched RNA sequencing, including both deep and low-pass WGS samples. Then, we randomly assigned the branch lengths to low-pass WGS samples, and repeated this procedure 100 times to obtain a median estimate of phylogenetic signal, as described in detail by ref^21^. Mean neoantigen/non-antigenic expression was computed by taking the average of the values log(x+1), where x is the expression (in TPM) of a gene within the subset evaluated. In both cases, we limited the analysis to cancer samples and to cases that had at least 6 biopsies in the final pruned trees.

### Analysis of H&E slides

Sections were cut from diagnostic formalin-fixed-paraffin-embedded blocks sampled at resection. H&E slides pre- and post-LCM and post-CyCIF were scanned using the NanoZoomer S210 slide scanner (Hamamatsu, Welwyn Garden City, UK). Representative sections were selected for annotations, which were drawn on pre-LCM H&E slides as first choice in most cases. Annotations included:

- all tumour (on a slide),
- the tumour-normal interface (which was later expanded by 500μm on either side to identify the invasive margin,
- any deposits of cancer within nodes,
- adjacent normal mucosa (normal_adjacent: within 5mm of superficial tumour), and normal mucosa further away (normal_far: >5mm away from tumour).

Annotations were made using the NDPViewer software (Hamamatsu, Welwyn Garden City, UK). The tumour-normal interface was expanded by 500μm on either side to identify the invasive margin. Images with annotations were analysed using a digital cell classifier which uses deep learning methodology modelled on a spatially constrained neural network architecture^29^. The model first detects cells via predicted location of cell nuclei, then classifies them as: normal epithelial, cancer epithelial, fibroblast, lymphocyte, neutrophil, macrophage, endothelial. Absolute counts are calculated for each annotated region. For each annotation, number of cells of a given type per epithelial cell was identified by dividing total number of cells of that type by total number of epithelial cells for that annotation.

In addition to the above, we used a further classifier^35^ to class all tumour-associated lymphocytes as infiltrating (ITL) or adjacent (ATL) or distant (DTL) based on proximity to tumour cells.

### CyCIF imaging

ROIs that matched the cancer glands/ micro-biopsies (enriched for epithelial cells) used for laser capture microdissection (LCM) were identified using the H&Es pre-LCM and post-LCM and extended by an additional 1mm at the edges. All CyCIF sections were 5 μm thick. The maximum separation between the LCM slides and the CyCIF slide was 10μm.

The protocol was based on previous methods from Lin et al.^22^. Full details of fluorescent antibodies are listed in Supplementary Table S1.

### Analysis of CyCIF images

One raw image file was acquired in the .ndpi format per cycle of scanning. Raw images were cropped using ImageJ^46^ such that all images were centred on the region of interest (ROI) that was serial to that used for genomics at LCM. Images were cropped to the same width and height dimensions across cycles to enable registration. Differences in background illumination were corrected using the built-in rolling ball algorithm in ImageJ.

The MCMICRO pipeline^47^ was used for registration (ASHLAR), segmentation (UnMICST/ S3segmenter) and signal quantification (MCQuant). Autofluorescence was removed by subtracting the intensity values noted for FITC and TRITC during the background scan (with no markers) from intensity values for markers in the first 3 marker cycles using R. Artefacts were identified by plotting the coordinates of cells showing expression values in the 99.95^th^ quantile for each marker on the raw image (i.e. the cells with the highest expression) using R. If such cells were localised to an illumination artefact, then those cells were removed from all further analysis by specifying coordinates for the affected area.

For each sample, for each marker, coordinates of cells with expression above a range of quantiles were plotted serially on the raw image (as red dots) for that marker using R. Images were then reviewed manually: a quantile where red dots overlapped with most positive cells on the raw image with few false positives was selected as the quantile threshold for that marker. All markers for all samples were reviewed similarly to identify a quantile threshold. Any values lower than the quantile threshold were set to 0.0001 for each marker. After thresholding, for each marker, expression per cell was divided by the 99.99th quantile for that marker using R. This allowed us to scale expression for different markers to similar ranges to allow us to identify co-expression of markers using clustering.

Using R, coordinates of all cells identified using MCMICRO were plotted on a raw DAPI mask. The “clickpoly” function from the R package spatstat v2.3-4 was used to interactively draw a polygon around the ROI. The polygon was expanded by a small amount of buffer region (50 units) to minimise any differences across ROIs in manually drawing the boundary. Only cells falling within the ROI were subsetted for further analysis.

The final list of 20 markers used for clustering was: Ki67, iNOS, CD45, CD8, IDO1, PD1, CD163, CD3, PDL1, CD4, CD68, VISTA, CD20, CTLA4, CD45RO, MYELO, ECAD, FOXP3, VIMENTIN, CK.

Expression of all markers for all cells from all CyCIF ROIs (1627807 cells in total) were clustered with R PhenoGraph, the R implementation for the PhenoGraph algorithm (R package: Rphenograph v0.99.1) with k=45. PhenoGraph takes as input a matrix of N single-cell measurements and partitions them into subpopulations by clustering a graph that represents their phenotypic similarity^48^. 262 clusters were identified. For each cluster, median normalised expression of every marker was identified. For a given cluster, if median normalised expression was greater than 0 for a marker, the cluster was considered positive for that marker.

32 clusters with no positive markers were excluded from further analysis. Such clusters may represent cells with signal below the threshold applied in previous steps for a marker (with each such cell showing marker expression 0.0001). 3 clusters showing positive markers that would not be expected together – i.e. a mixture of epithelial/ stromal/ immune markers – were also excluded. Otherwise, the following rules were used for labelling phenotypes, implemented in R:

Epithelial cells:

- Only CK/ECAD/ both: phenotype of clusters was labelled as “epithelial cells”.
- CK/ECAD/ both with Ki67: phenotype of clusters was labelled as “proliferating epithelial cells”.
- Clusters expressing ECAD/CK/both and Vimentin were labelled “Mixed epithelial/stromal” and excluded from further analysis.

Stromal cells:

- Only Vimentin: phenotype of clusters was labelled as “stromal cells”.

Immune cells:

- If only 1 immune marker was included, the phenotype was labelled as the marker followed by a “+” sign.
- Multiple markers:

`o` If 2 or more markers were present with all being immune markers, all makers were included in the final phenotype.
`o` If 2 or more markers were present with at least one immune marker and a second marker being “ECAD/CK/ Vimentin /CD45”, this additional second marker was ignored in the final phenotype. This was because:

i. An assumption was made that mixed epithelial/ stromal component with immune markers was due to difficulties cleanly segmenting infiltrating immune cells from epithelial/ stromal background. In this situation, the immune marker was more relevant to the phenotype than the contaminating epithelial/stromal component.
ii. As CD45 is the lymphocyte common antigen, it does not add additional distinguishing subtype-specific information. For example, clusters which are CD45+CD3+ and CD3+ only both show a phenotype of T cells (the subtype-specific marker is paramount in this case and both clusters would be labelled CD3+ for analysis purposes). Similarly, cluster 116 (showing CD45,CD8,CD3, cluster 254 showing CD8,ECAD,CK and 162 showing CD8 only were labelled as “cytotoxic T cells”.
iii. Clusters expressing only CD45 or CD45 with epithelial/stromal component were labelled as “Lymphocyte, not otherwise specified”.
`o` The rule under 4b for phenotyping of immune/epithelial clusters was not applied to PDL1 and CK/ ECAD as PDL1 may be expressed by epithelial or immune cells. Cluster 120 expressing PDL1 CK and 122 expressing PDL1 ECAD were both classified as “PDL1+ epithelial cells”.

After applying the above rules, 29 phenotypes were identified in total. 3 phenotypes were excluded for further analysis (“NA”, “Mixed epithelial/ stromal”, “PD-1+ MYELO+”), leaving 26 phenotypes for downstream analysis.

#### Visual assessment of phenotypes

For each ROI, coordinates of a subset of cells were plotted over the raw DAPI image used for segmentation to visually assess appropriateness of phenotyping. Phenotypes which were prevalent across ROIs such as epithelial cells, stromal cells, lymphocyte (no specific type) and cytotoxic T cells were selected for visual assessment. All epithelial cell phenotypes (“Epithelial cells”, “PDL1+ epithelial cells” and “Proliferating epithelial cells”) were grouped together as “Epithelial cells”. Additionally, the commonly prevalent immune markers of CD3/CD4/CD68/CD163/CD20/MYELO were selected such that any cells labelled with a cluster positive for any one of these markers would be included.

Where cells were labelled for more than one included marker, the marker pertaining to the broadest category of cells was used as the final label. For example, if a cell was positive for both CD3 and CD4, CD3 was prioritised as the label. Similarly, CD68 was prioritised over CD163. This was to allow easy visualisation of broad categories rather than niche subtypes.

### Statistical analysis

All statistical analysis was carried out in R (version 4.x). Unless stated otherwise, all comparisons between sample groups (e.g. stratified by immune escape status or sample type) were carried out using unpaired Wilcoxon rank sum test, without additional adjustment of p-values.

Paired Wilcoxon rank sum test was used to compare values derived from the same sample/patient in Figures XX as indicated by lines connecting paired observations.

#### Evaluating neoantigens in expressed genes and not expressed neoantigens

We created 2×2 contingency tables by counting mutations that are neoantigens/non-antigenic and (i) in a gene in the given gene group or not; (ii) in a consistently expressed gene or not; (iii) not expressed (missing) in at least one sample or not. Fisher’s exact test was used to compute the Odds ratio and confidence intervals on these tables. The above steps were repeated separately on clonal and subclonal mutations alone.

#### Distance between epithelial and immune cells

The R package spatstat v2.3-4 was used to determine the pairwise Euclidean distances between cells of different phenotypes. For each ROI, cells of the two phenotypes of interest were identified and pairwise Euclidean distances established using the function crossdist(). For each cell of interest, the closest cell of the second phenotype was determined by taking the shortest pairwise distance of all cells of the second phenotype. The Euclidean distance in pixels was converted to micrometres by multiplying the distance by 0.44μm, the size of each pixel.

For this analysis, phenotypes “proliferating epithelial cells” and “epithelial” cells were considered together as epithelial cells (PDL1+ epithelial cells were not included here). All phenotypes that were positive for Ki67+ in addition to a cell type marker were clubbed together. In other words, phenotypes “Ki67+CD20+ and “CD20” were clubbed together, as were “Ki67+MYELO+” / “MYELO+” and “Ki67+PDL1+” / “PDL1+”. Where all lymphocytes were considered together, they were grouped as described above.

#### Multivariable regression

To test the dependence of proportional burden and immune dNdS on other variables, we constructed multivariable regression models using the functions *betareg* (for proportional burden) and *lm* (for immune dNdS). Sample type (gland vs bulk for FF-WGS and superficial tumour/invasive margin/node for FFPE-PS), tissue type (adenoma vs cancer), immune escape (escape/weak escape/no escape) and patient were encoded as categorical variables; and purity as a continuous variable. Results were visualised using the plot_summs function from the package jtools, omitting coefficients assigned to patients to make the visuals easier to interpret.

#### Ripley’s H Index

Ripley’s H-index^49^ was calculated using the Tumour Landscape Analysis pipeline (https://github.com/cisluis/TLA/blob/main/documentation/TLA_doc.md). For each reference cell, the number of test cells I_rt_(d) inside a radius d from the reference cell is established. The Ripley’s K function is the mean across all reference cells normalized by the density λ_t_ of test cells (overall density across the entire ROI).

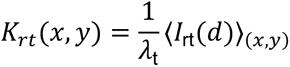

*K* is effectively the proportion of observed to expected points in the circle surrounding a reference cell of radius d. When the distribution of test points is homogeneous (i.e. number of test cells within the circle is similar to number expected based on overall test cell density), (i) the expected value of I should approach A λ_t_ with A= πd^2^ (the area of the circle); (ii) the expected value of K should approach A (with A= πd^2^).

The *H* function is defined as:

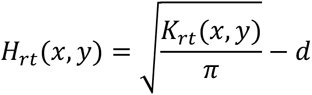

The H function is a measure of the level of clustering of test cells around reference cells at the scale *d*. When the number of test points within the circle is similar to number expected based on overall test cell density, K= πd^2^, H=0. When the number of test points within the circle is lower or higher than expected, H becomes negative or positive, respectively. *H* ~ 0 indicates that reference and test cells are mixed uniformly. We used d=100 μm as interactions between immune cells and epithelial cells are likely to occur within this proximity – this metric has been used in previous work from our collaborators^165^.

#### Cellular neighbourhoods (CN)

We adapted the method of cellular neighbourhood classification from Schürch et al.^36^. For each of the 1,146,536 cells across all ROIs, a ‘window’ was captured consisting of the 10 nearest neighbouring cells (including the centre cell) as measured by Euclidean distance between X/Y coordinates. This was an unsupervised approach implemented through NearestNeighbors (from sklearn.neighbours) with the “auto” algorithm (default). These windows were then clustered by their composition with respect to the 26 cell types that had previously been using Python’s *scikit-learn* implementation of MiniBatchKMeans with number of neighbourhoods *k* = 15. In this manner, 15 cellular neighbourhoods were identified, which were each enriched for a variety of cell types. Each cell was then allocated to the CN that its surrounding window was in. To validate the CN assignment, these allocations were overlaid on the original nuclear mask from the fluorescent images.

For each CN, for a cell type, the mean number of nearest neighbours matching this cell type represented in the CN was noted (“cluster “centroids”). For every cell type, the proportion of cells in this cell type out of all cells (from all ROIs) was established. This proportion was added to the cluster centroids for the matching cell type. The overall sum was divided by 11 (the sum of mean number of nearest neighbours (10 for each cell) and sum of proportions for all cell types (1) and was then divided by the proportion of cells for a cell type (as determined above) to normalise for cell type frequency. Based on the cell type composition for each CN, each CN was given a unique label. Every cell across all ROIs was assigned to a CN.

To compare immune cell subtypes within CNs, the 6 CNs positive for epithelial cells were identified, namely CN1: “Epithelial cell-enriched”, CN0: “Tumour-immune interface I”, CN5: “Tumour-immune interface II”, CN14: “Tumour-immune interface III”, CN3: “PDL1+ and PDL1-epithelial cell-enriched” and CN10: “Cells expressing ICRs, CD4+ and macrophage-enriched”. CNs positive for PDL1+ epithelial cells (CN3 and CN10) were compared to the other CNs. Immune cell types were defined as all phenotypes other than “Epithelial cells”, “Proliferating epithelial cells”, “PDL1+ epithelial cells”, “stromal cells” and “Ki67+ only”.

To compare specific cell types within CNs across sample types, the fraction and number per epithelial cell for each cell type of interest (e.g. PD1+, PD1+CD3+CD45RO+, PDL1+ and PDL1+ epithelial cells) within each CN in each ROI were identified and compared measures across sample types.

## References

1. Hanahan, D. Hallmarks of Cancer: New Dimensions. Cancer Discov. 12, 31–46 (2022).

2. Varn, F. S., Wang, Y., Mullins, D. W., Fiering, S. & Cheng, C. Systematic pan-cancer analysis reveals immune cell interactions in the tumor microenvironment. Cancer Res. 77, 1271–1282 (2017).

3. Vilar, E. & Gruber, S. B. Microsatellite instability in colorectal cancer—the stable evidence. Nat. Rev. Clin. Oncol. 7, 153–162 (2010).

4. Alex J. Cornish et al. Whole genome sequencing of 2,023 colorectal cancers reveals mutational landscapes, new driver genes and immune interactions. bioRxiv 2022.11.16.515599 (2022) doi:10.1101/2022.11.16.515599.

5. Alouani, E., et al. Efficacy of immunotherapy in mismatch repair-deficient advanced colorectal cancer in routine clinical practice. An AGEO study. ESMO Open 8, (2023).

6. Le, D. T. et al. PD-1 Blockade in Tumors with Mismatch-Repair Deficiency. N. Engl. J. Med. 372, 2509–2520 (2015).

7. Pagès, F. et al. International validation of the consensus Immunoscore for the classification of colon cancer: a prognostic and accuracy study. Lancet Lond. Engl. 391, 2128–2139 (2018).

8. Lakatos, E. et al. Evolutionary dynamics of neoantigens in growing tumors. Nat. Genet. 52, 1057–1066 (2020).

9. Newey, A. et al. Immunopeptidomics of colorectal cancer organoids reveals a sparse HLA class I neoantigen landscape and no increase in neoantigens with interferon or MEK-inhibitor treatment. J. Immunother. Cancer 7, 309 (2019).

10. van den Bulk, J., et al. Neoantigen-specific immunity in low mutation burden colorectal cancers of the consensus molecular subtype 4. Genome Med. 11, 87 (2019).

11. Giannakis, M. et al. Genomic Correlates of Immune-Cell Infiltrates in Colorectal Carcinoma. Cell Rep. 15, 857–865 (2016).

12. Angelova, M. et al. Evolution of Metastases in Space and Time under Immune Selection. Cell 175, 751–765.e16 (2018).

13. Grasso, C. S. et al. Genetic Mechanisms of Immune Evasion in Colorectal Cancer. Cancer Discov. 8, 730–749 (2018).

14. Gatenbee, C. D. et al. Immunosuppressive niche engineering at the onset of human colorectal cancer. Nat. Commun. 13, 1798 (2022).

15. Heide, T. et al. The co-evolution of the genome and epigenome in colorectal cancer. Nature 611, 733–743 (2022).

16. Zhu, G. et al. ARID1A, ARID1B, and ARID2 Mutations Serve as Potential Biomarkers for Immune Checkpoint Blockade in Patients With Non-Small Cell Lung Cancer. Front. Immunol. 12, 670040 (2021).

17. Corces, M. R. et al. The chromatin accessibility landscape of primary human cancers. Science 362, eaav1898 (2018).

18. Rosenthal, R. et al. Neoantigen-directed immune escape in lung cancer evolution. Nature 567, 479–485 (2019).

19. Gai, X.-D., Li, C., Song, Y., Lei, Y.-M. & Yang, B.-X. In situ analysis of FOXP3+ regulatory T cells and myeloid dendritic cells in human colorectal cancer tissue and tumor-draining lymph node. Biomed. Rep. 1, 207–212 (2013).

20. Bindea, G. et al. Spatiotemporal dynamics of intratumoral immune cells reveal the immune landscape in human cancer. Immunity 39, 782–795 (2013).

21. Househam, J. et al. Phenotypic plasticity and genetic control in colorectal cancer evolution. Nature 611, 744–753 (2022).

22. Lin, J.-R. et al. Highly multiplexed immunofluorescence imaging of human tissues and tumors using t-CyCIF and conventional optical microscopes. eLife 7, e31657 (2018).

23. Zapata, L. et al. Immune selection determines tumor antigenicity and influences response to checkpoint inhibitors. Nat. Genet. 55, 451–460 (2023).

24. Schenck, R. O., Lakatos, E., Gatenbee, C., Graham, T. A. & Anderson, A. R. A. NeoPredPipe: high-throughput neoantigen prediction and recognition potential pipeline. BMC Bioinformatics 20, 264 (2019).

25. Pagel, M. Inferring the historical patterns of biological evolution. Nature 401, 877–884 (1999).

26. Freckleton, R. P., Harvey, P. H. & Pagel, M. Phylogenetic analysis and comparative data: a test and review of evidence. Am. Nat. 160, 712–726 (2002).

27. Khanna, A. et al. Bam-readcount - rapid generation of basepair-resolution sequence metrics. J. Open Source Softw. 7, 3722 (2022).

28. Angelova, M. et al. Characterization of the immunophenotypes and antigenomes of colorectal cancers reveals distinct tumor escape mechanisms and novel targets for immunotherapy. Genome Biol. 16, 64 (2015).

29. Sirinukunwattana, K. et al. A Spatially Constrained Deep Learning Framework for Detection of Epithelial Tumor Nuclei in Cancer Histology Images. in Patch-Based Techniques in Medical Imaging (eds. Wu, G., Coupé, P., Zhan, Y., Munsell, B. & Rueckert, D.) 154–162 (Springer International Publishing, Cham, 2015). doi:10.1007/978-3-319-28194-0_19.

30. Challoner, B. R. et al. Genetic and immune landscape evolution defines subtypes of MMR deficient colorectal cancer. 2022.02.16.479224 Preprint at 10.1101/2022.02.16.479224 (2022).

31. Jongsma, M. L. M., Guarda, G. & Spaapen, R. M. The regulatory network behind MHC class I expression. Mol. Immunol. 113, 16–21 (2019).

32. McGranahan, N. et al. Allele-Specific HLA Loss and Immune Escape in Lung Cancer Evolution. Cell 171, 1259–1271.e11 (2017).

33. Martínez-Jiménez, F. et al. Genetic immune escape landscape in primary and metastatic cancer. Nat. Genet. 55, 820–831 (2023).

34. Yuan, L., Tatineni, J., Mahoney, K. M. & Freeman, G. J. VISTA: A Mediator of Quiescence and a Promising Target in Cancer Immunotherapy. Trends Immunol. 42, 209–227 (2021).

35. Yuan, Y. Modelling the spatial heterogeneity and molecular correlates of lymphocytic infiltration in triple-negative breast cancer. J. R. Soc. Interface 12, 20141153 (2015).

36. Schürch, C. M. et al. Coordinated Cellular Neighborhoods Orchestrate Antitumoral Immunity at the Colorectal Cancer Invasive Front. Cell 182, 1341–1359.e19 (2020).

37. McGranahan, N. et al. Clonal neoantigens elicit T cell immunoreactivity and sensitivity to immune checkpoint blockade. Science 351, 1463–1469 (2016).

38. Wolf, Y. et al. UVB-Induced Tumor Heterogeneity Diminishes Immune Response in Melanoma. Cell 179, 219–235.e21 (2019).

39. Westcott, P. M. K. et al. Low neoantigen expression and poor T-cell priming underlie early immune escape in colorectal cancer. *Nat*. Cancer 2, 1071–1085 (2021).

40. Chowell, D. et al. Improved prediction of immune checkpoint blockade efficacy across multiple cancer types. Nat. Biotechnol. 40, 499–506 (2022).

41. Bruni, D., Angell, H. K. & Galon, J. The immune contexture and Immunoscore in cancer prognosis and therapeutic efficacy. Nat. Rev. Cancer 20, 662–680 (2020).

42. Sottoriva, A. et al. A Big Bang model of human colorectal tumor growth. Nat. Genet. 47, 209–216 (2015).

43. Shukla, S. A. et al. Comprehensive analysis of cancer-associated somatic mutations in class I HLA genes. Nat. Biotechnol. 33, 1152–1158 (2015).

44. Kim, S. et al. Strelka2: fast and accurate calling of germline and somatic variants. Nat. Methods 15, 591–594 (2018).

45. Favero, F. et al. Sequenza: allele-specific copy number and mutation profiles from tumor sequencing data. Ann. Oncol. 26, 64–70 (2015).

46. Fiji: an open-source platform for biological-image analysis | Nature Methods. https://www.nature.com/articles/nmeth.2019.

47. Schapiro, D. et al. MCMICRO: a scalable, modular image-processing pipeline for multiplexed tissue imaging. Nat. Methods 19, 311–315 (2022).

48. Levine, J. H. et al. Data-Driven Phenotypic Dissection of AML Reveals Progenitor-like Cells that Correlate with Prognosis. Cell 162, 184–197 (2015).

49. Ripley, B. D. Modelling Spatial Patterns. J. R. Stat. Soc. Ser. B Methodol. 39, 172–192 (1977).

