## Supplementary Table and Figures for "Epigenome and early selection determine the tumour-immune evolutionary trajectory of colorectal cancer"

| Cycle | FITC (green) | Target cell type | Dilution | TRITC (yellow) | Target cell type | Dilution | Cy5 (red) | Target cell type | Dilution |
|---|---|---|---|---|---|---|---|---|---|
| 1 | Ki67 | Proliferation | 200 | iNOS | M1 macrophages | 100 | CD45 | All white blood cells | 50 |
| 2 | CD8 | Cytotoxic T cells | 100 | IDO1 | Immune checkpoint (antigen presenting cells) | 100 | PD1 | Immune checkpoint (T cells) | 50 |
| 3 | CD163 | M2 Macrophages | 200 | CD3 | Pan T cells | 100 | PDL1 | Immune checkpoint (epithelial)- occasional T cells can be positive | 50 |
| 4 | CD4 | Helper T cells | 50 | CD68 | Macrophages | 200 | Vista | Immune checkpoint (T cells) | 50 |
| 5 | CD20 | B cells | 100 | CTLA4 | Immune checkpoint (T cells) | 100 | CD57 | NK Cells | 100 |
| 6 | CD45RO | Memory T cells | 100 | HLA-ABC | HLA expression | 600 | Myeloperoxidase | Neutrophils | 400 |
| 7 | E-cadherin | Epithelial cells | 100 | FOXP3 | Regulatory T cells | 100 | Vimentin | Mesenchymal cells | 3000 |
| 8 | N/A | N/A | N/A | Pan CK | Epithelium | 400 | N/A | N/A | N/A |

**Supplementary Table 1. Panel of markers used in each CyCIF cycle.**

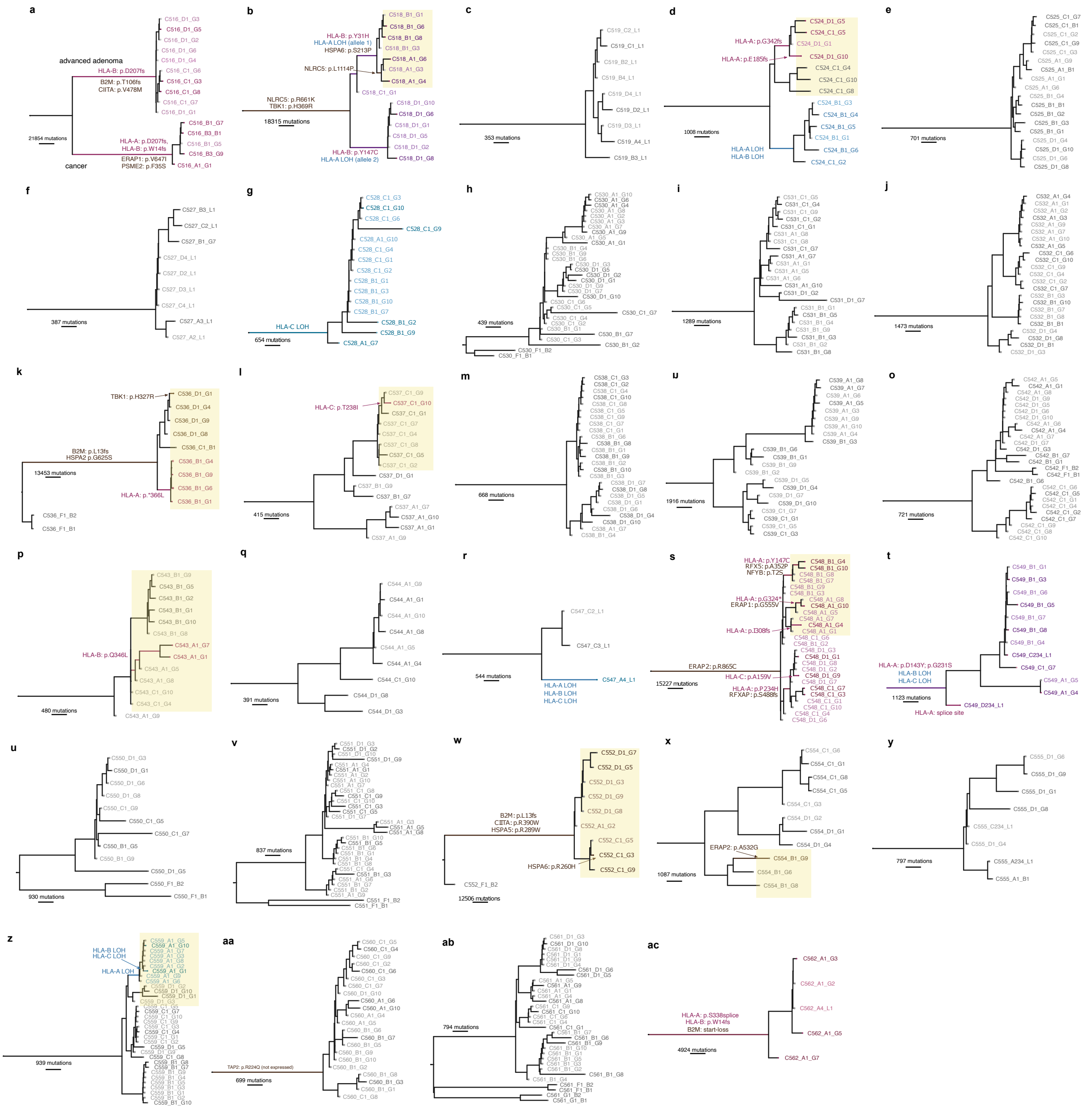


**Supplementary Figure 1. Phylogenetic trees reconstructed using FF-WGS data of the 29 CRCs.** Immune escape alterations are shown by arrows or over the branch they occurred in. Light-coloured samples were sequenced using low-pass WGS and genotyped using deep-sequenced samples. Biopsies in yellow shaded areas in (b,d,k,l,p,s,w,x & z) are subclones considered for subclone-specific normalised burden and immune dNdS analysis in Fig. 5c-d.

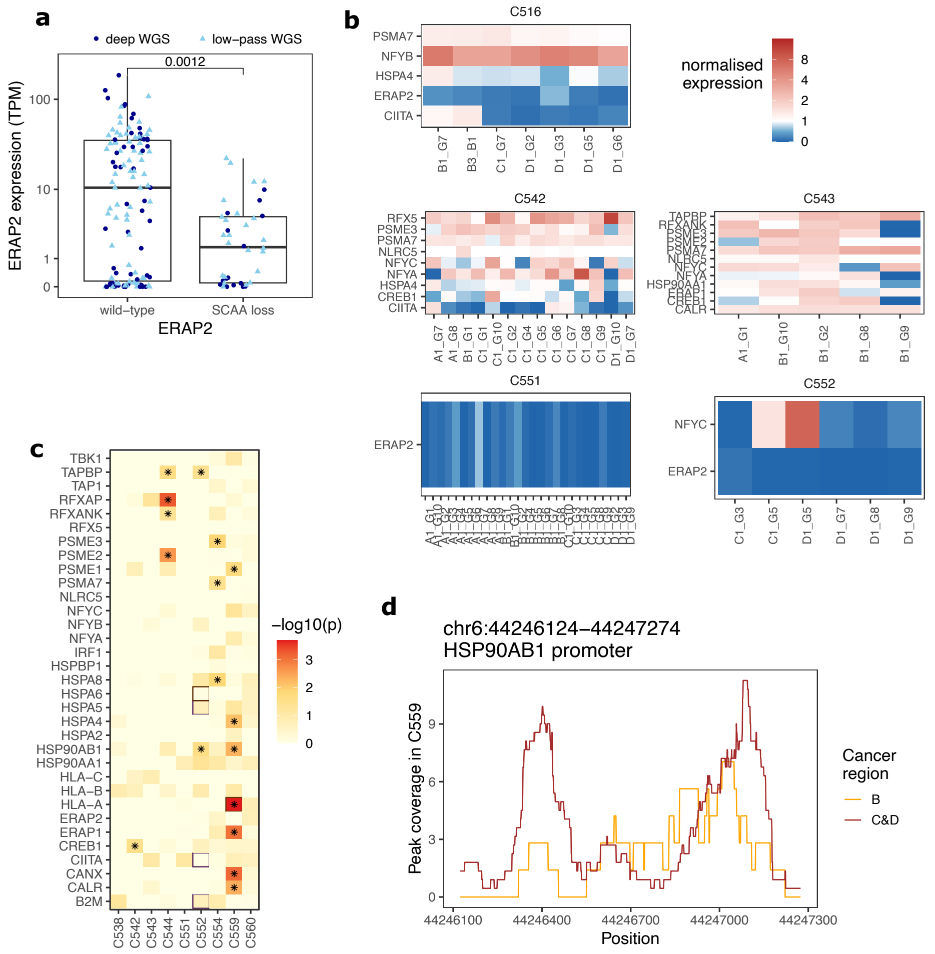


**Supplementary Figure 2. Chromatin-associated and heritable regulation of antigen presenting genes**. (a) Expression of ERAP2 in samples with and without a SCAAloss in the promoter of the gene. (b) Normalised expression (compared to non-tumour samples) of antigen presenting genes affected by SCAAloss. (c) Phylogenetic signal of the expression of genes associated with immune escape. Cancers where expression of a particular gene is phylogenetic ($p<0.05$) are indicated by asterisks. Cancer-gene combinations with a somatic mutation of the gene are indicated by brown rectangles. (d) Peak coverage of the HSP90AB1 promoter region in cancer C559. Maroon and orange lines show the average across all samples within region C&D and B, respectively.

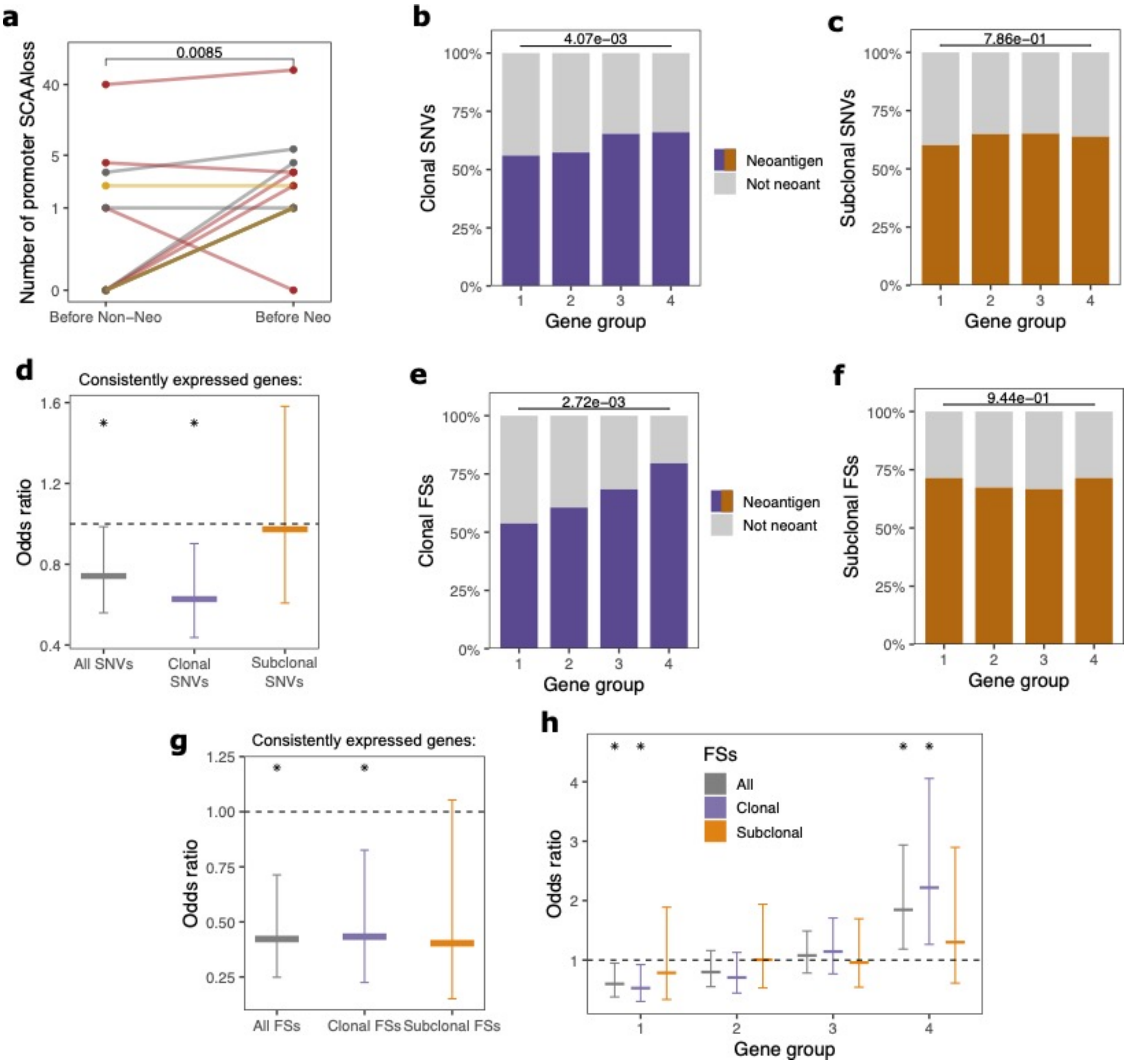


**Supplementary Figure 3. Epigenetic and transcriptomic immuno-editing** (a) Total number of SCAAlosses falling in promoters of genes with an antigenic or non-antigenic mutation. (b-c,e-f) The proportion of antigenic (in colour) and non-antigenic (in grey) SNVs (b,c) and FSs (e,f) that are located in genes of gene group 1-4 from ref[31]. Clonal (b,e) and subclonal (c,f) mutation are shown separately. The p-value of a chi-squared test comparing groups is shown on top of each panel. (d,g) Odds ratio of a neoantigen vs non-antigenic SNVs (d) and FSs (g) being located in consistently expressed genes. (h) Odds ratio of a neoantigen vs non-antigenic FS mutation being located in genes of gene group 1-4 from ref[31]. All, clonal and subclonal mutations are shown separately in grey, purple and orange. In d&g-h markers represent OR values, and error bars show confidence intervals.

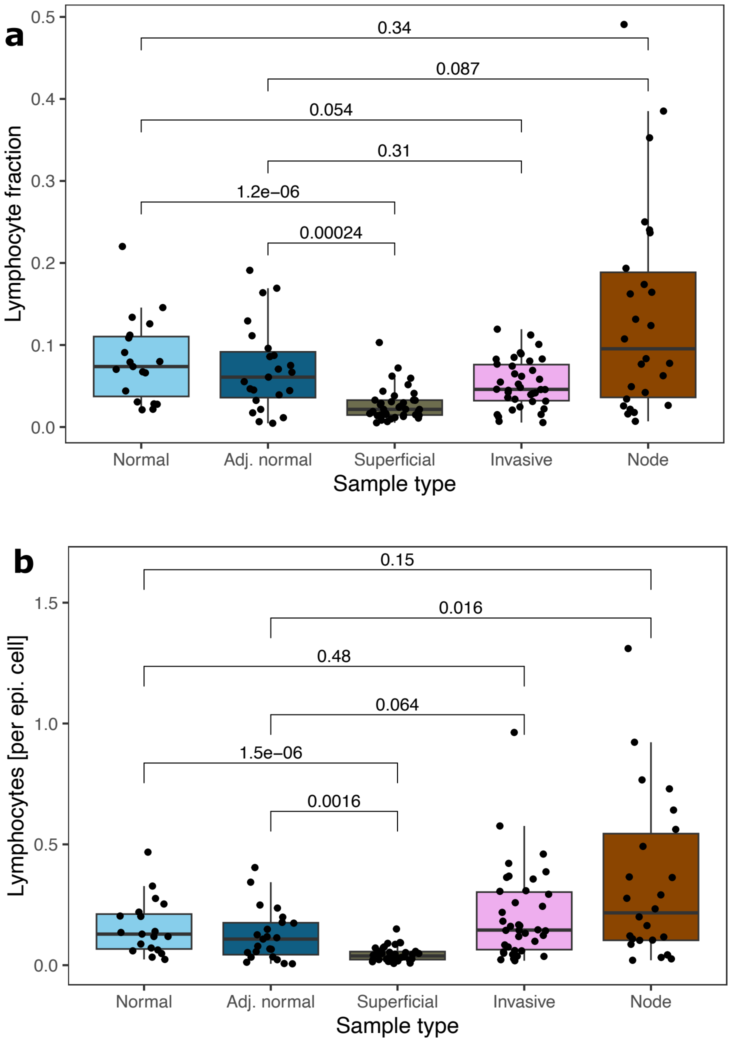


**Supplementary Figure 4. Lymphocyte infiltration quantified from H&E images.** (a,b) Fraction of lymphocytes (a) and number of lymphocytes normalised by epithelial cell count (b) in normal mucosa and tumour-containing regions. Lymphocyte counts were derived from H&E stained images using a deep learning-based cell classifier.

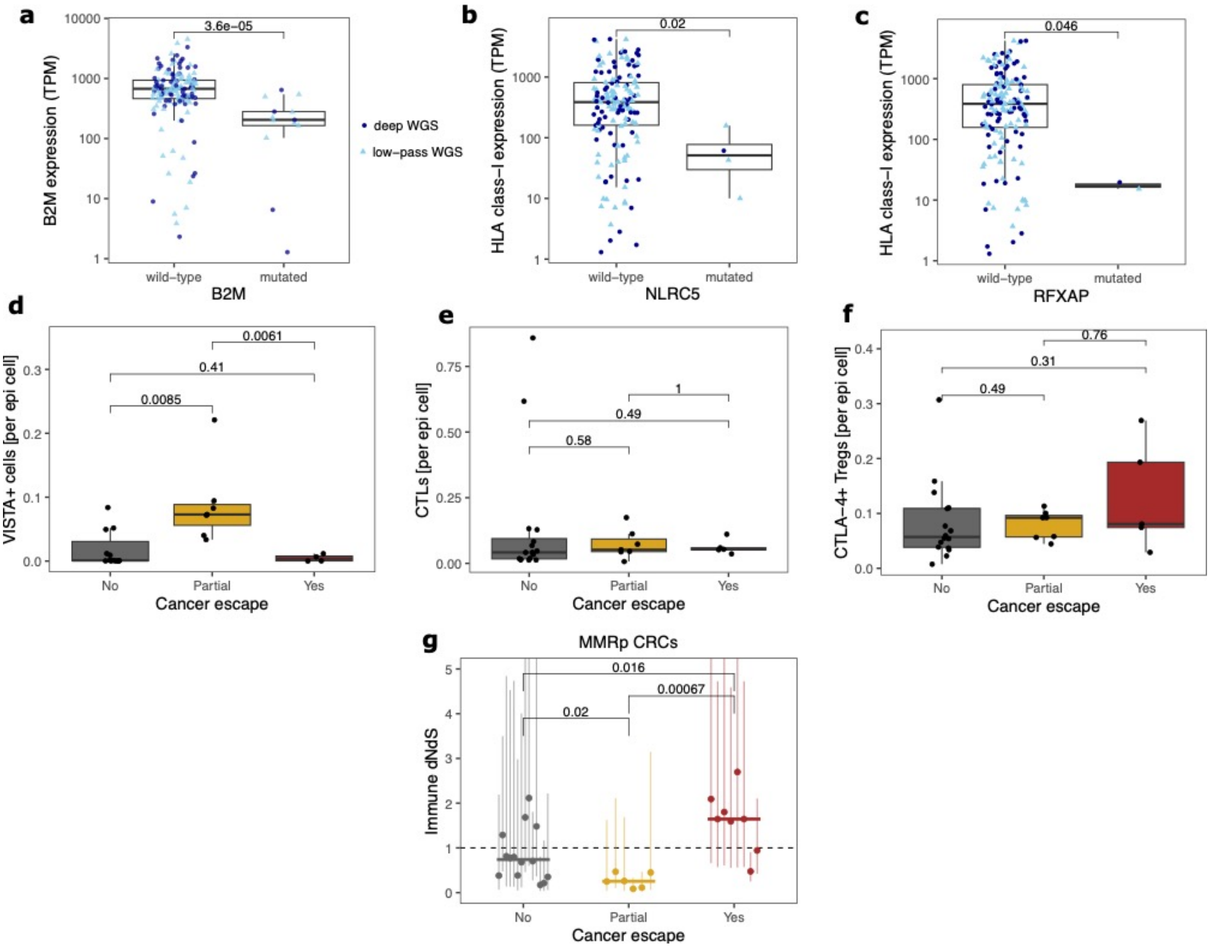


**Supplementary Figure 5. Relationship between immune escape mutations and immune measures.** (a) Expression of B2M in samples with and without a mutation in the gene. (b,c) Total combined expression of HLA class-I genes (HLA-A, -B and -C) compared in samples with and without a mutation in NLRC5 (b) and RFXAP (c). (d-f) Number of VISTA+ cells (d), CTLs (e) and CTLA-4+ Tregs (f) per epithelial cell, shown based on cancer immune escape status. (g) Immune dNdS values in MMRp FF-WGS cancers. Each dot represents a biopsy, and error bars show confidence intervals of immune dNdS values

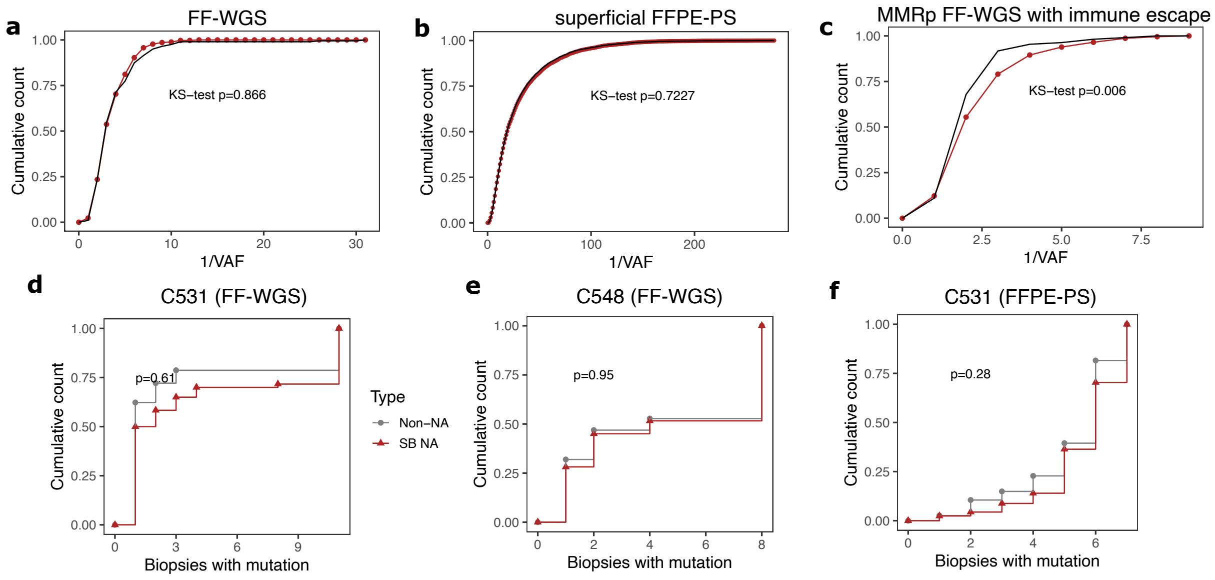


**Supplementary Figure 6. Frequency distribution of neoantigens**. (a-c) Cumulative number of non-antigenic (grey) and neoantigen (red) mutations shown against the inverse of the variant allele frequency. All mutations were pulled together from FF-WGS non-escaped cancers (a), FFPE-PS superficial tumour samples (b), or FF-WGS escaped MMRp samples. (d-f) The cumulative distribution of the number of mutations shared by a given number of biopsies, shown for non-neoantigen (in grey) and neoantigen (in red) mutations in the indicated cancers. Kolmogorov-Smirnov test results are shown above the graphs.

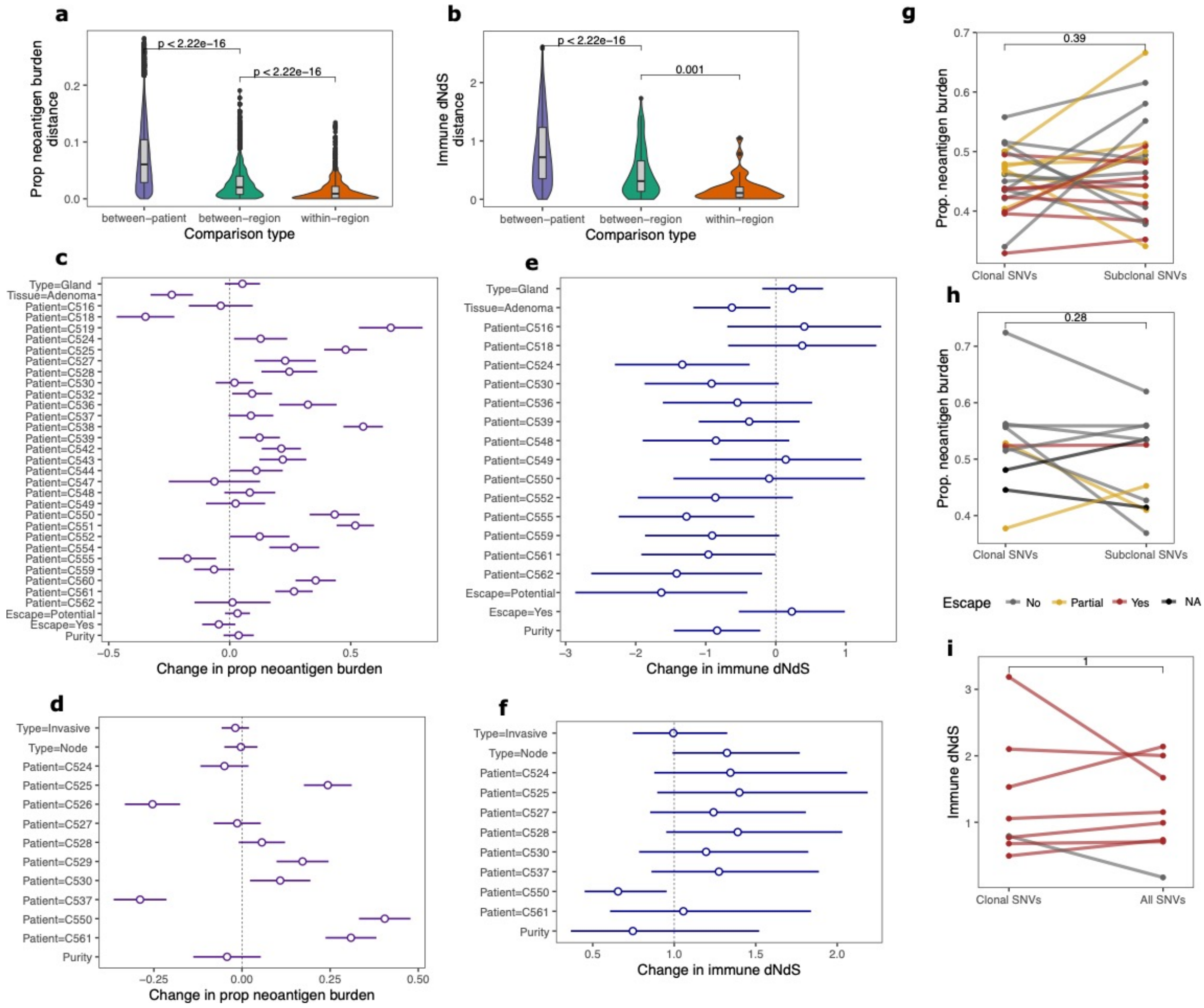

**Supplementary Figure 7. Intra-tumour heterogeneity of neoantigen landscape and immuno-editing in CRCs.** (a,b) Distribution of pairwise differences in proportional neoantigen burden (a) and immune dNdS (b) between FF-WGS biopsies from the same tumour region (orange), from different tumour regions of the same tumour (green) and from different cancers (purple). (c-f) Forest plots depicting the output of a multivariable regression (Methods) investigating the association of proportional neoantigen burden (c,d) and immune dNdS (e,f) with other sample characteristics, in FF-WGS (c,e) and FFPE-PS (d,f). Circles denote the estimated coefficients, with whiskers showing 95% confidence intervals. (g-h) Comparison of clonal and subclonal neoantigen burden in FF-WGS cancers (g) and in FFPE-PS cancers (h). Each line indicates paired values derived from the same cancer. (i) Comparison of immune dNdS computed from clonal and all mutations in FF-WGS cancers. Each line indicates paired values derived from the same cancer.

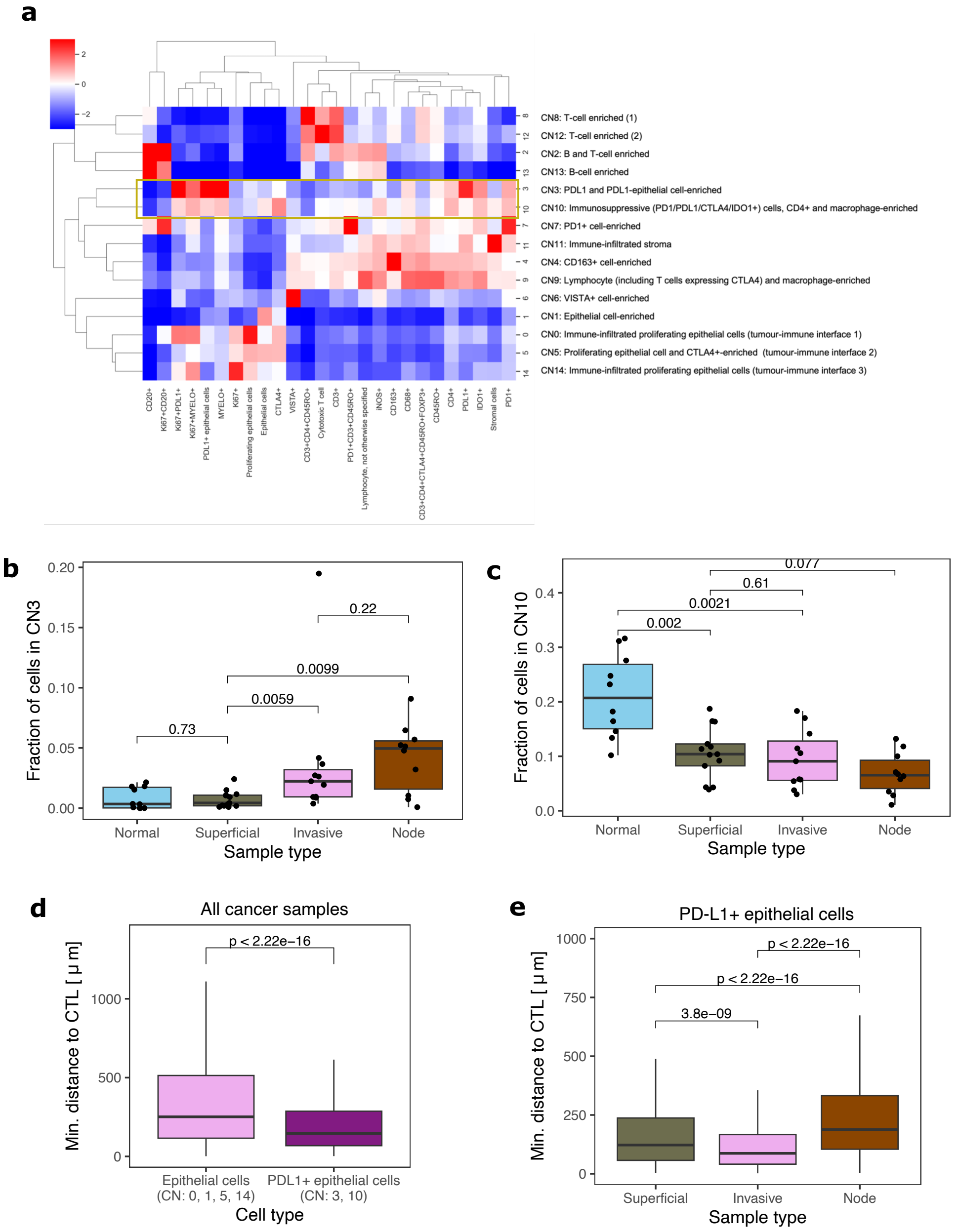


**Supplementary Figure 8. PD-L1+ cellular neighbourhoods and cell-cell interactions.** (a) Heatmap depicting the derived cellular neighbourhoods and the relative abundance of cell types in each neighbourhood. Rows and columns are ordered according to hierarchical clustering. CN3 and CN10 are highlighted by the yellow frame. (b-c) Fraction of cells in each sample type that belong to cellular neighbourhoods 3 (b) and 10 (c). (d) Distance of epithelial cells to the closest cytotoxic T-cell, for non-PDL1-expressing epithelial cells in non-PD-L1+ CNs and for PD-L1+ epithelial cells in PD-L1-associated CNs, in all cancer samples. (e) Distance of PD-L1+ epithelial cells (from CN3 and 10) to the closest cytotoxic T-cell, shown separately for tumour sample types.

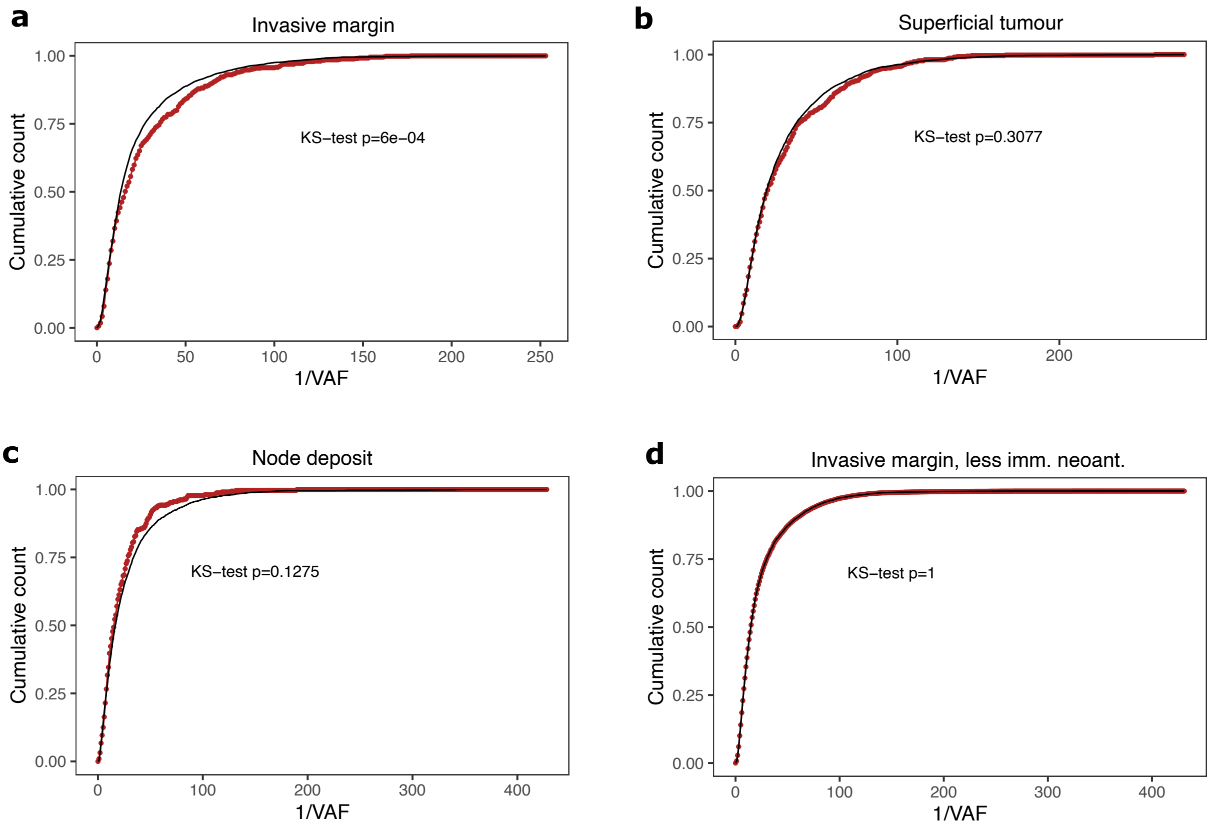


**Supplementary Figure 9. Frequency distribution of neoantigens in different tumour sample types.** The cumulative number of low-immunogenicity mutations (grey) and high-immunogenicity neoantigens (red) shown against the inverse of the variant allele frequency. All mutations were pulled together from FFPE-PS samples from the invasive margin (a,d), superficial tumour (b) or node (d). In (a-c), immunogenic neoantigens are defined as strong-binders with high recognition potential and compared to weak-binders with low recognition potential. In (d), they are defined as strong-binders and compared to non-binders.
